# Distributed activity in the human posterior putamen distinguishes goal-directed from habitual control

**DOI:** 10.1101/2025.11.26.690863

**Authors:** Amogh Johri, Lisa Kluen, Rani Gera, Vincent Man, Omar Perez, Julia Pia Simon, Weilun Ding, Aniek Fransen, Scarlet Cho, Sarah Oh, Jeff Cockburn, Jamie D. Feusner, Reza Tadayonnejad, John P. O’Doherty

## Abstract

How do some individuals rapidly form habits while others maintain flexible, goal-directed control? Using multivariate fMRI decoding in 199 participants, we show that distributed neural activity patterns in the left posterior putamen during initial learning predict individual behavioral strategy on a subsequent outcome devaluation test. This prediction generalized across two independent cohorts of healthy adults and psychiatric patients with heterogeneous diagnoses and was anatomically specific to the posterior putamen. Critically, predictive neural signatures were present during training, before strategy expression after devaluation, enabling prospective classification of habitual versus goal-directed behavior. These findings demonstrate that stable individual differences in behavioral control are reflected in circumscribed brain activity during learning, highlighting the posterior putamen as a candidate neural marker of habit propensity with potential clinical relevance.

## Introduction

Reward-driven action selection in both humans and non-human animals relies on at least two distinct modes of behavioral control: goal-directed and habitual strategies [1–3]. Goal-directed control involves the deliberate selection of actions based on the current value of expected outcomes, allowing for flexible adaptation when goals or circumstances change. In contrast, habitual control is reflexive and insensitive to the value of the action’s outcome.

Considerable research has characterized the neural mechanisms of goal-directed and habitual control in both humans and other animals. Brain regions, including the ventromedial prefrontal cortex, dorsolateral prefrontal cortex, anterior dorsal striatum, and putative homologues in the rodent brain, like the prelimbic cortex and the dorsomedial striatum, have been implicated in goal-directed control [4–6]. Conversely, the posterior putamen and its rodent equivalent, the dorsolateral striatum, have been implicated in habitual control [7–11]. In humans, the posterior putamen activity increases with training [9, 12] and encodes multivariate representations of stimuli and actions, but not outcomes. In contrast, the dorsolateral and ventromedial prefrontal cortex encode representations of actions, outcomes, and goals [13].

However, individuals vary remarkably in the behavioral expression of goal-directed and habitual control [14]. Some individuals form specific habits rapidly, while others take much longer, or may not form them at all within a given timeframe [15]. Understanding the neural basis of these individual differences has both theoretical and clinical significance. Maladaptive habits underlie addiction and compulsive disorders [16–18], yet we cannot predict who will develop rigid, outcome-insensitive behavior. Hence, a critical unresolved question is whether brain activity during initial learning can prospectively predict whether a given individual will subsequently adopt a habitual or goal-directed behavioral strategy.

To address this question, participants performed a free-operant action selection task during fMRI, in which sensitivity to outcome devaluation indexed habitual versus goal-directed control (Figure 1). We used multivariate pattern analysis to decode behavioral strategy from neural activity during the training phase, before devaluation. Leveraging a large fMRI sample [19], we implemented a fully held-out validation framework, training classifiers in an initial cohort and testing generalization in independent samples of healthy participants and psychiatric patients with heterogeneous diagnoses. This design enabled a stringent test of whether neural activity during learning prospectively predicts individual behavioral strategy across populations.

We implemented all analyses in Python and provide fully reproducible results using publicly available code (see Data and materials availability).

**Fig. 1:**
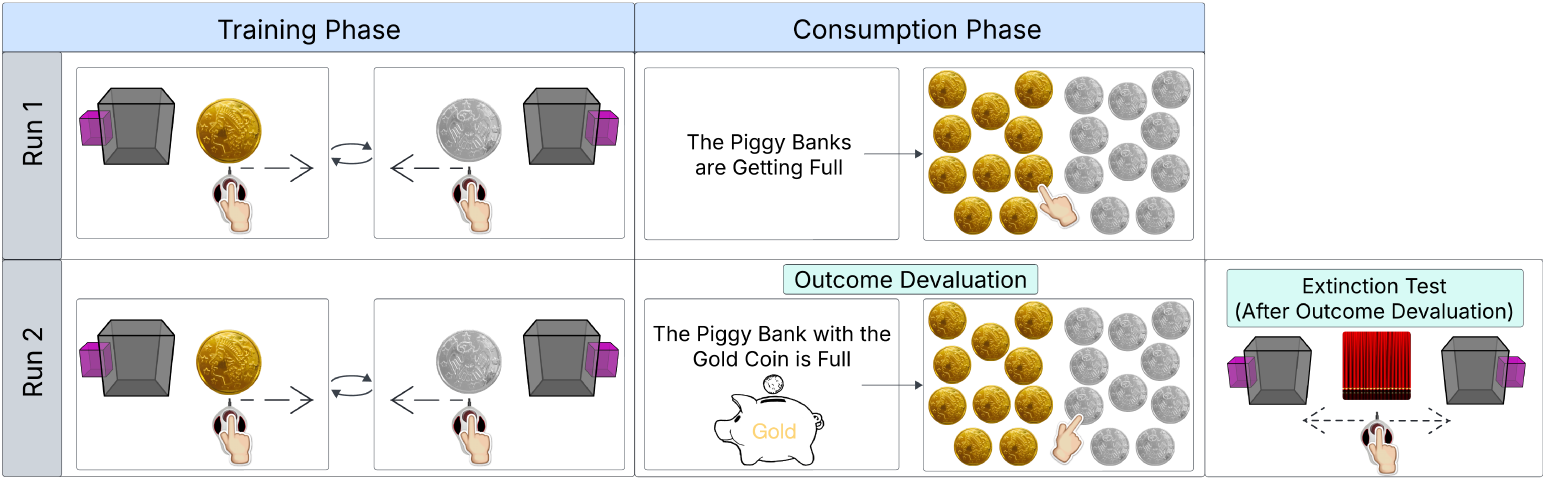
Free-operant outcome-devaluation task design for dissociating goal-directed and habitual control. *Pre-task instructions (not shown).* Before scanning, participants were instructed that they would earn gold and silver coins of equal value by performing actions in response to visual stimuli, and that coins earned during the task would be accumulated in separate virtual piggy banks and later exchanged for monetary reward. *Training phase.* Participants learned stimulus–action–outcome associations during free-operant training, with each block involving a single stimulus–outcome pairing (gold or silver). Coins were collected via leftward or rightward swiping on a trackball according to a random-ratio schedule. The mapping between stimuli, actions, and outcomes was fully counterbalanced across participants and remained fixed within individuals. Training comprised 10 alternating blocks per run (20 total; 37–60 s per block). *Consumption phase.* Following training, participants completed a consumption phase to assess sensitivity to outcome devaluation. Participants collected a fixed number of coins in the absence of stimuli. Outcome devaluation was administered only during the second run by informing participants that one of the piggy banks was full. *Extinction test.* Participants were presented with both stimuli simultaneously, and participants freely selected which action to perform under pseudo-extinction conditions, such that actions no longer produced visible rewards. Preferential selection of the action associated with the non-devalued outcome, indexed to goal-directed control, whereas continued responding for the devalued outcome, indexed to habitual control. Further task details are in the Methods (Task Description).

## Results

We investigated whether neural activity during the training phase of a free-operant action-selection task predicted whether individuals subsequently expressed goal-directed or habitual behavior following outcome devaluation. We split the healthy sample (*n* = 144) into an Initial Cohort (75%*, n* = 108) and Held-Out Cohort (25%*, n* = 36) before all analysis. We trained a binary classifier on an initial cohort of healthy participants using cross-validation, and evaluated out-of-sample generalization on the Held-Out Cohort. In addition to the analysis, we further evaluated whether the neural signature identified in healthy participants generalized to a separate Held-Out Patient Cohort (*n* = 55) with heterogeneous psychiatric diagnoses.

### Behavioral Results

#### Devaluation Ratio

We computed a devaluation-ratio for each participant, defined as the proportion of actions (swipes) directed toward the still-valued coin relative to the total number of actions during the choice test (see Methods (Task: Choice Test Phase)). A devaluation-ratio of 1.0 indicates an exclusive response associated with the still-valued outcome, consistent with perfectly goal-directed behavior. In contrast, a devaluation-ratio of 0.5 reflects an equal preference for response in both the devalued and still-valued outcomes, consistent with habitual responding. As illustrated in Figure 2A, participants exhibited a broad spectrum of behavior, with peaks at goal-directed (devaluation-ratio = 1.0) and habitual (devaluation-ratio = 0.5).

Although the devaluation ratio is continuous in form, its interpretability is regime-dependent. In the habitual regime, variation largely reflects stochastic responding under pseudo-extinction, whereas in the goal-directed regime it reflects meaningful sensitivity to outcome devaluation. Accordingly, devaluation ratios were strongly bimodal, motivating categorical stratification for primary analyses.

To facilitate subsequent analyses, we categorized participants into goal-directed or habitual based on their devaluation ratios using a data-driven threshold derived from a two-component Beta mixture model (see Methods (Group Classification)). This model identified two latent subpopulations corresponding to distinct behavioral strategies and estimated a posterior decision boundary at a devaluation-ratio of 0.648. Participants with devaluation ratios at or above this threshold were classified as goal-directed, reflecting a predominant preference for the still-valued outcome. In contrast, participants with devaluation ratios below the threshold were classified as habitual, reflecting outcome-insensitive responding. This classification divided the Initial Cohort (*n* = 108) into two comparably sized groups (Goal-Directed: *n* = 55; Habitual: *n* = 53), providing a principled basis for subsequent binary subgroup analyses of neural and behavioral differences.

#### Learning Related Changes in Response Rates

Participants completed 10 training blocks in each of two runs. For each run, we fit a linear mixed-effects regression model on participants from the Initial Cohort, with block number, group, and their interaction as fixed effects, and subject as a random intercept (see Methods: Response Rates).

**Fig. 2:**
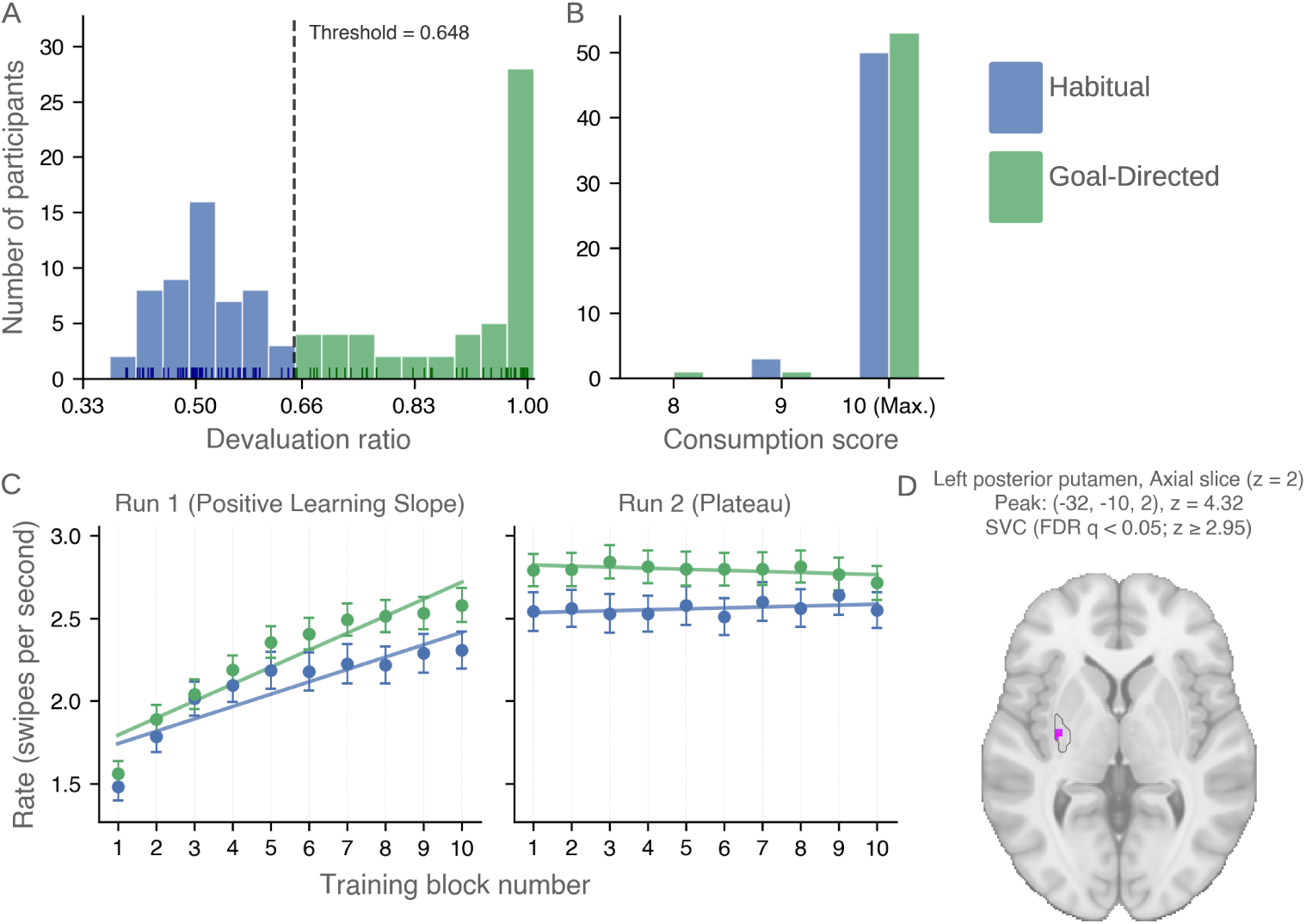
Behavioral stratification, learning dynamics, and neural correlates during training. All plots include participants after exclusions (see Methods (Exclusions)) from the Initial Cohort (*n* = 108). **(A)** Distribution of devaluation ratios across participants. Values near 1.0 indicate preferential selection of the still-valued outcome (goal-directed behavior), whereas values near 0.5 indicate indifference (habitual behavior). The devaluation-ratio threshold of 0.648 (dashed line) divided the Initial Cohort (*n* = 108) into Goal-Directed (*n* = 55) and Habitual groups. **(B)** Distribution of consumption scores following outcome devaluation. Scores are discrete (8–10), with the majority of participants in both groups achieving the maximum score, indicating near-ceiling performance. **(C)** Mean response rates (swipes/s) across training blocks for Habitual and Goal-Directed participants, shown separately for Run 1 and Run 2. Points denote group means, and error bars indicate ±1 standard error of the mean (SEM); solid lines represent fitted linear trends. **(D)** Learning-related fMRI signal in the left posterior putamen. An axial slice (z = 2) shows voxels exhibiting increasing signal across training blocks, identified using a parametric regressor modeling linear learning-related change. Results are small-volume corrected within the left posterior putamen (FDR *q <* 0.05; *z* ≥ 2.95). The peak voxel is located at (−32, −10, 2) (z = 4.32).

During Run 1, response rates increased significantly across blocks, consistent with within-session learning (Figure 2C). The model revealed a robust main effect of block number, indicating increasing response rates over time. Notably, there was a significant group × block interaction: goal-directed participants exhibited a steeper increase in response rate (0.112 swipes/s/block, *z* = 15.75*, p <* 0.001) than habitual participants (0.077 swipes/s/block, *z* = 6.16*, p <* 0.001). The difference in slopes was statistically significant (*β* = −0.036*, z* = −3.52*, p <* 0.001), indicating greater learning-related increases in responding among goal-directed participants.

In Run 2, response rates showed no significant within-group change across blocks (goal-directed: −0.0096 swipes/s/block, *z* = −1.62*, p* = 0.105; habitual: 0.0072 swipes/s/block, *z* = 0.70*, p* = 0.484), consistent with a plateau in performance. However, a small but significant group × block interaction was observed (*β* = 0.017*, z* = 1.99*, p* = 0.047), reflecting diverging but weak trends across groups (habitual slightly increasing; goal-directed slightly decreasing). In addition, there was a significant group difference in intercept (*β* = −0.564*, z* = −2.74*, p* = 0.006), with goal-directed participants maintaining higher overall response rates throughout Run 2.

Together, these results indicate that learning occurs primarily during the first run, after which performance stabilizes. Goal-directed participants exhibit a steeper learning-related increase in Run 1 and maintain an overall advantage in response vigor across both runs compared to habitual participants.

#### Behavioral and Trait-Related Predictors of Devaluation Sensitivity

Next, we examined behavioral and cognitive predictors of devaluation sensitivity within the Initial Cohort using a logistic regression predicting goal-directed classification (threshold = 0.648; 1 = goal-directed) from standardized predictors (KNN-imputed missing values, *k* = 5; Type-III Wald tests, see Methods (Model Specification)). The model converged (pseudo-*R*^2^ = 0.230; LLR *p* = 0.043). Higher response rates during training were associated with a greater likelihood of goal-directed behavior (Wald *χ*^2^(1) = 4.546, *p* = 0.033; *β* = 0.599).

To assess explicit contingency knowledge, participants completed post-task probes of response–outcome associations by indicating the action required to obtain each outcome (a silver or gold coin; a left or right swipe) (see Methods (Task: Contingency Memory Assessment)). Greater response–outcome knowledge was significantly associated with goal-directed behavior (Wald *χ*^2^(1) = 8.737, *p* = 0.003; *β* = 0.869). In contrast, stimulus–response knowledge, assessed by presenting the initial cues and asking participants to select the corresponding action, was not significantly associated with habitual responding (Wald *χ*^2^(1) = 0.144, *p* = 0.704; *β* = −0.084).

When extending this analysis to the full sample (*n* = 199; healthy controls and patients), the predictive effect of response rate attenuates to a non-significant trend. In addition, we find higher OCI-R scores associated with increased habitual behavior (Wald *χ*^2^(1) = 4.359, *p* = 0.036; *β* = −0.467), consistent with prior reports linking compulsivity to reduced goal-directed control (Supplementary: Figure S3).

#### Training-Related Changes in fMRI Activity

Next, we examined learning-related changes in fMRI activity during training by modeling the signal across training blocks (see Methods (fMRI Analysis)). We fit a general linear model (GLM) to fMRI data from the Initial Cohort (*n* = 108), including a parametric modulator that increased linearly with block number to capture learning-related trends. Brain regions positively associated with this regressor were interpreted as exhibiting increasing activity across training.

This analysis revealed a significant cluster in the left posterior putamen (Figure 2D), where BOLD activity increased progressively over training blocks. Results were small-volume corrected within the anatomically defined region of interest (ROI; see Methods (ROIs)) using false discovery rate correction (FDR *q <* 0.05; cluster-size threshold = 10 voxels). The analysis was restricted a priori to striatal ROI, and we didn’t test on whole-brain or non-striatal ROIs.

#### Neural Decoding of Goal-Directed vs. Habitual Control

A central aim was to identify distributed neural activity patterns that distinguish individuals who exhibit predominantly goal-directed versus predominantly habitual control. To address this, we implemented a subject-wise multivariate decoding approach to classify participants based on their neural responses during instrumental training.

We evaluated neural decoding of goal-directed versus habitual control within the Initial Cohort (*n* = 108) using four-fold cross-validation. Neural features were derived from a block-wise general linear model (GLM) fit separately for each participant (see Methods (First Level)). All training blocks (10 per run; 20 total) were modeled with block regressors and contrasted with the implicit baseline, yielding a single-subject-level beta map capturing multivoxel activation patterns during instrumental training.

To mitigate the confounding effects of individual differences in motor output, we regressed each participant’s mean response rate and response-rate slopes (separately for each run) from the neuroimaging data before decoding. All subsequent analyses were performed on the resulting residuals (see Methods (Confound Regression)).

We trained a Linear Discriminant Analysis (LDA) classifier to predict group membership (goal-directed vs. habitual). LDA provides a linear, low-variance classifier suitable for high-dimensional neuroimaging data and offers a conservative test of multivoxel discriminability. Classification performance was quantified using mean cross-validated accuracy; accuracy is appropriate here given the near-equal group sizes (55 vs. 53).

We assessed statistical significance using non-parametric permutation testing with 5,000 random label shuffles for each voxel-centered searchlight. We perform inference using permutation-based small-volume family-wise error correction within each ROI, with an additional Bonferroni correction applied across the 11 predefined ROIs (see Methods (MVPA Statistical Analysis)).

#### Primary Decoding Result on Initial Cohort

Our primary decoding analysis was restricted to a priori ROIs previously implicated in goal-directed and habitual control [13], including the posterior putamen, anterior caudate, dorsolateral prefrontal cortex (dlPFC), and ventromedial prefrontal cortex (vmPFC). Within this anatomically constrained search space, we identified a single cluster in the left posterior putamen in which decoding accuracy was reliably above permutation-derived null (mean cross-validated accuracy = 0.627 ± 0.023 SD; 60 voxels; small-volume family-wise error corrected *p* = 0.0198, with Bonferroni correction across 11 ROIs; Figure 3A).

The left-lateralized selectivity in the posterior putamen may reflect the strong right-handed bias in our sample (97% right-handed in the Initial Cohort), consistent with contralateral motor representation in the basal ganglia. However, the present analysis does not directly test for hemispheric asymmetries and cannot rule out alternative explanations for this lateralization.

To assess decoding effects outside the predefined ROIs, we additionally performed a whole-brain searchlight analysis. No regions outside the a priori ROIs showed statistically reliable classification accuracy after correction for multiple comparisons, highlighting the anatomical specificity of posterior putamen involvement in distinguishing goal-directed from habitual individuals (Supplementary: Anatomical specificity across decoding approaches).

#### Generalization to an Independent Held-Out Cohort

Next, we tested whether the posterior putamen decoder identified in the Initial Cohort would generalize to unseen participants using a Held-Out Cohort (*n* = 36, 25% of healthy participants). We classified participants as goal-directed or habitual using the same devaluation-ratio threshold (0.648), defined a priori based solely on the Initial Cohort. The threshold yielded 13 habitual and 23 goal-directed individuals in the Held-Out Cohort (Figure4A).

We applied a region-based decoder restricted to the 60-voxel cluster in the left posterior putamen identified by the searchlight analysis. To approximate the effective spatial resolution of the searchlight spheres and enhance generalizability, the decoder was implemented as a bagging ensemble of linear discriminant classifiers (see Methods (Generalization Analysis)), each trained on random feature subsets. This design reduces variance across ensemble members and mirrors the feature subsampling inherent to the searchlight procedure.

As the Held-Out Cohort was somewhat imbalanced (13 habitual vs. 23 goal-directed participants), we assessed decoder performance using ROC AUC, a threshold-independent measure of separability that is more appropriate than raw accuracy under class imbalance. Re-evaluating performance in the Initial Cohort with this 60-voxel ROI yielded a mean ROC AUC of 0.656 ± 0.018 SD (Figure3C). We present the permutation-based null distribution for reference; however, we do not report inferential statistics for the Initial Cohort at this stage, as such tests would be dependent on the ROI selection procedure.

**Fig. 3:**
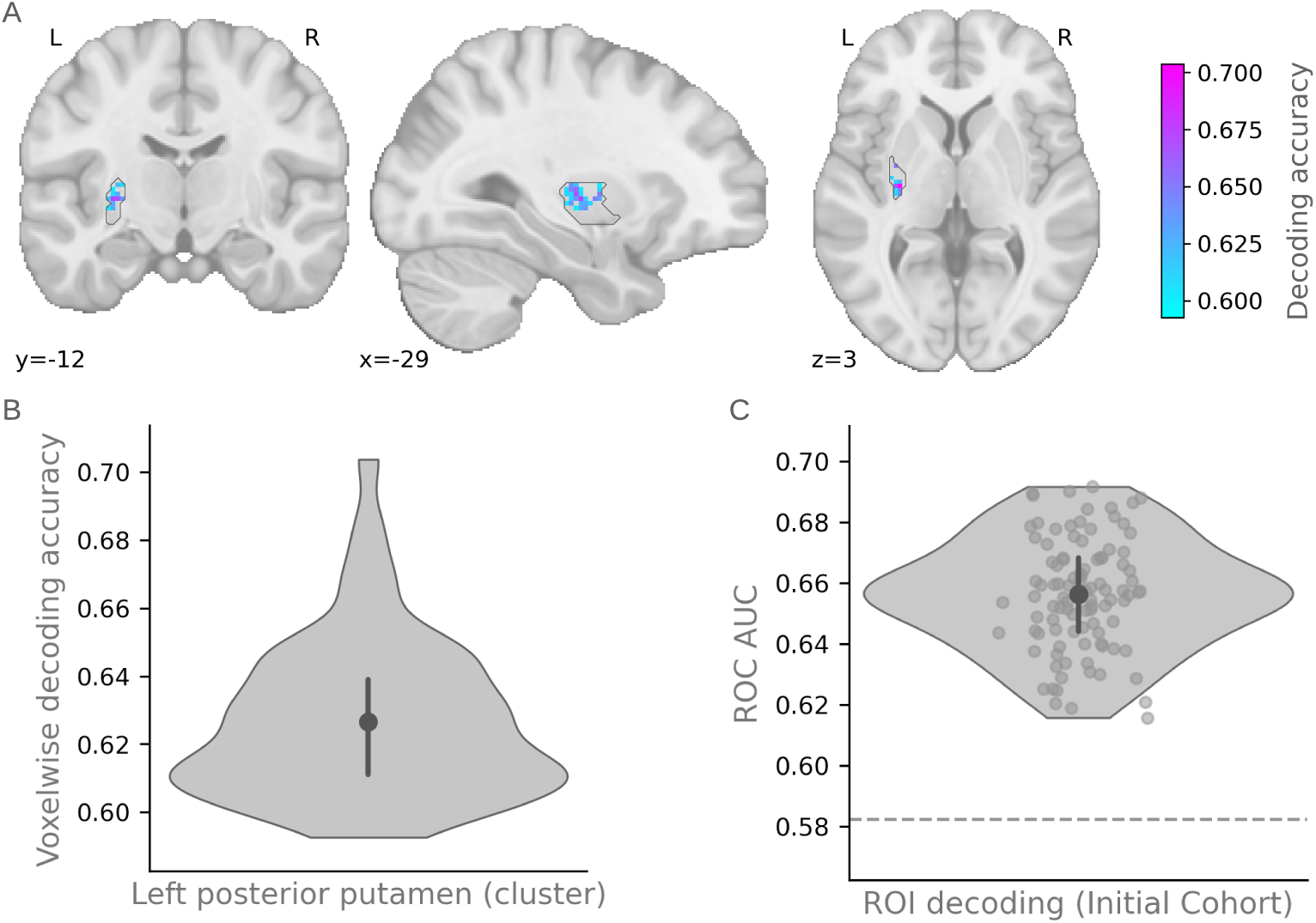
Multivariate decoding of habitual versus goal-directed behavioral strategy in the left posterior putamen. **(A)** Voxel-wise map showing mean cross-validated decoding accuracy within the significant left posterior putamen cluster, displayed on the MNI template (coronal: y = −12; sagittal: x = −29; axial: z = 3). **(B)** Distribution of voxel-wise mean cross-validated decoding accuracies within the left posterior putamen cluster. The violin plot shows the distribution of accuracy values across voxels. The central point indicates the mean accuracy (0.627), and the vertical bar represents the interquartile range (25th–75th percentile). **(C)** ROI-level decoding performance treating the entire significant left posterior putamen cluster as a single region. The violin plot shows the distribution of ROC AUC values across repeated cross-validation estimates. The central point indicates the mean ROC AUC (0.656), and the vertical bar represents the interquartile range (25th–75th percentile). The dashed horizontal line indicates the 95th percentile of the permutation-derived null distribution.

Crucially, the Held-Out Cohort was completely untouched during model training: the decoder was trained exclusively on the Initial Cohort and then applied directly to the Held-Out Cohort without any retraining or parameter tuning. Applying it to the Held-Out Cohort yielded a mean ROC AUC of 0.666 ± 0.020 SD, significantly above permutation-derived null based on 5,000 label permutations (*p* = 0.044; Figure 4B). These results demonstrate that posterior putamen activity reliably distinguishes habitual from goal-directed participants in an independent sample. As detailed in Supplementary: Table S4, performance was robust across alternative decoder implementations, indicating that the observed effect is not dependent on a specific classifier choice.

**Fig. 4:**
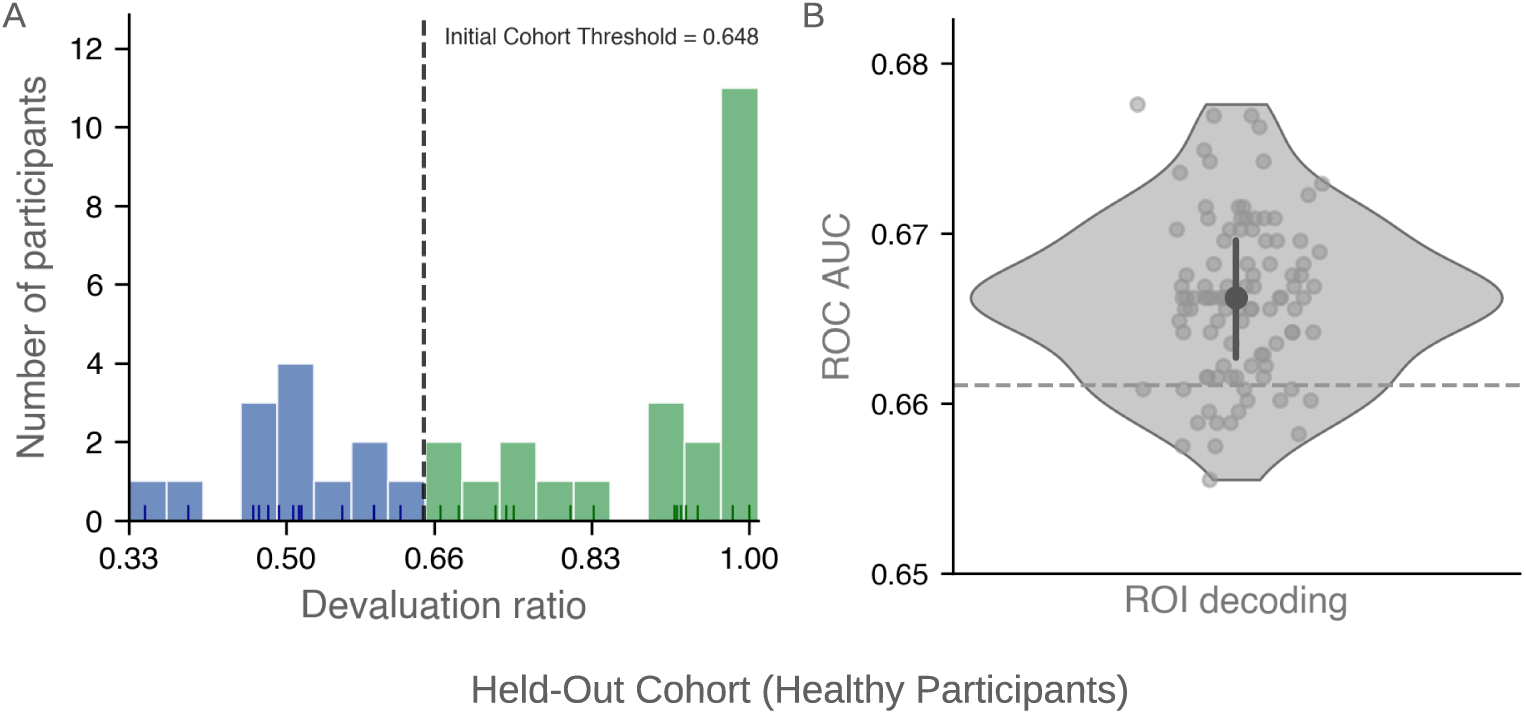
Generalization of habitual versus goal-directed classification and neural decoding to a held-out cohort. **(A)** Distribution of devaluation ratios in the Held-Out Cohort (healthy participants, *n* = 36), classified using the same devaluation ratio threshold (0.648) defined a priori in the Initial Cohort. Green bars indicate goal-directed participants; blue bars indicate habitual participants. **(B)** ROI-level decoding performance in the Held-Out Cohort. The decoder, trained exclusively on the Initial Cohort, was applied to the held-out data without retraining or parameter tuning. The violin plot shows the distribution of ROC AUC values across repeated decoder evaluations. The central point indicates the mean ROC AUC (0.666), and the vertical bar represents the interquartile range (25th–75th percentile). Performance was significantly above the permutation-derived null (*p* = 0.0442), indicated by the dashed horizontal line (95th percentile).

#### Generalization to an Independent Psychiatric Patient Cohort

Our sample also included a cohort of psychiatric patients with heterogeneous diagnoses (N = 55 after exclusions; see Supplementary Table S1 and Table S2). We tested whether behavioral strategy could be decoded in this independent held-out population using the same devaluation-ratio threshold defined in the Initial Cohort of healthy participants, which divided the patient cohort into 26 goal-directed and 29 habitual individuals. Applying the significant posterior putamen cluster from the Initial Cohort, the decoder achieved robust separability in this sample (ROC AUC = 0.739 ± 0.019 SD), significantly above the permutation-derived null (*p* = 0.001). These results indicate that both the behavioral classification and its associated posterior putamen signal generalize across diagnostically diverse populations, providing further replication of the decoding effect (see Supplementary: Figure S4).

#### Nature of Representations (Multivariate vs. Univariate)

We asked whether the posterior putamen decoding effect reflects multivariate pattern information, univariate signal differences, or both. First, we regressed out the mean (univariate) signal from each participant’s voxel pattern within the MVPA cluster. Decoding performance remained significantly above permutation-derived null (ROC AUC = 0.609, *p* = 0.022), indicating that multivariate pattern structure alone carries discriminative information.

We then examined whether univariate differences also contribute by averaging activation across all voxels within the cluster. Habitual participants exhibited numerically higher average activation than goal-directed participants (*t*(199) = 3.773, *p <* 0.001). This univariate effect did not survive voxel-wise small-volume correction across the broader posterior putamen ROI, but reached significance when aggregated across the MVPA-defined cluster.

We emphasize that these analyses are post hoc and exploratory, and the statistics are not independent inferential tests. Instead, they serve to characterize the nature of the signal, suggesting that the decoding effect reflects both distributed multivariate structure and modest, directionally consistent univariate differences within the cluster.

#### Additional robustness checks

We conducted supplementary analyses to assess whether decoding performance reflects stable neural structure rather than noise or analytic artifacts (see Supplementary: Robustness and Reproducibility of Multivariate Decoding and Supplementary: Graceful Degradation of Decoder Performance). Performance declined in a graded and interpretable manner when reducing the number of participants, subsampling voxels, increasing spatial smoothing, or filtering voxels by decoding-weight signal-to-noise ratio. Decoding performance remained stable across alternative cross-validation strategies and scoring metrics, and voxel-level weights exhibited consistent signs across cross-validation folds, with low variability across repetitions.

We further observed consistent results across alternative behavioral group-definition schemes, including the mixture-model boundary (devaluation-ratio = 0.648; AUC = 0.676 ± 0.010 SD), a median split (AUC = 0.679 ± 0.009 SD), and a top-versus-bottom quartile split (reduced *n*; AUC = 0.648 ± 0.020 SD; Supplementary: Table S7). We obtained comparable performance when decoding from the entire left posterior putamen ROI rather than the data-driven cluster (Supplementary: Figure S6), and when residualizing task-related, psychological, and demographic confounds before decoding (AUC = 0.629 ± 0.016 SD, *p <* 0.05).

Finally, a conventional MVPA searchlight analysis conducted on the full combined participant sample identified clusters in the left posterior putamen with high spatial overlap with the primary result (see Supplementary: Figure S5). Together, these analyses indicate that the decoding effect is robust across a wide range of analytic choices and is not strongly dependent on specific modeling decisions, cohort size, or residual confounds.

## Discussion

We find that distributed neural activity in the left posterior putamen during initial training of instrumental associations can distinguish between whether a given individual deploys a goal-directed or habitual strategy following a subsequent outcome devaluation procedure, a canonical test of goal-directed and habitual control in the animal learning literature [1, 20].

Critically, because the decoding procedure occurs during training rather than during the devaluation test itself, the decoder does not rely on trivial neural features associated with the devaluation test, such as reduced responding to the devalued outcome in the goal-directed group. Moreover, we controlled for confounding aspects of behavior during training that might distinguish goal-directed from habitual participants, including response rate and learning-curve slope. Notably, the participants classified as habitual did not merely misunderstand the devaluation procedure: the consumption test is a crucial element of our experiment that protects against this possibility. In this test, participants freely selected between the two token outcomes after devaluation but before the action-outcome devaluation test. Participants who do not understand the devaluation procedure would not show a substantial preference for the still-valued coin; we therefore excluded the small proportion who performed poorly on the consumption test. Thus, the participants in our sample provided clear evidence that they understood the devaluation procedure, as their coin preferences were consistent with optimal choice. Hence, this eliminates a confounding explanation: the classifier picks up on participants who exhibit habit-like behavior solely because of inattention to or misunderstanding of the devaluation procedure.

The decoder’s ability to generalize to an independent held-out sample provides strong evidence that neural patterns in the left posterior putamen reflect robust and behaviorally meaningful representations of individual differences in behavioral control strategy. Unlike most subject-wise MVPA studies that typically report only cross-validated performance within the same sample, our use of an ensemble classifier tested on unseen participants demonstrates true out-of-sample generalizability. It suggests that the identified multivoxel patterns do not overfit the training set but instead capture a consistent neural signature distinguishing goal-directed from habitual individuals across datasets. Importantly, the same classification approach also generalized to a cohort of patients with psychiatric diagnoses, indicating that this generalization extends beyond the original healthy participant sample.

The ability to distinguish behavioral control strategy from decoded posterior putamen activity suggests that neural activity in this region differs when individuals engage in goal-directed versus habitual behavior. This finding extends our understanding of the functional contributions of the posterior putamen while inviting several possible interpretations.

A longstanding proposal in the literature is that this region encodes stimulus-response associations that underpin learned habits [21–23]. It is possible that the goal-directed participants in the present study were less successful at learning stimulus-response associations, as reflected in different patterns of activity in the putamen during training compared to individuals who did learn robust stimulus-response associations and subsequently deployed habits, and that the decoder is exploiting these differences. It is relevant to note that we did not observe a significant relationship between stimulus-response knowledge (probed after the experiment) and goal-directedness, which would appear to indicate against this possibility. However, explicit knowledge of stimulus-response contingencies may not be synonymous with learned stimulus-response associations that are likely implicit and not easily accessible to meta-cognitive knowledge.

Alternatively, the posterior putamen may do more than merely encode stimulus-response associations, in that activity in this region could play an active role in implementing a specific mode of behavioral control. This possibility resonates with emerging evidence in the rodent literature indicating a relationship between neuronal activity in the rodent homologue of the posterior putamen, the dorsolateral striatum (DLS), and the behavioral expression of habits [24]. Previous studies have found neurons in DLS to decrease engagement during goal-directed actions [8], while DLS neuronal activity during the onset of a behavioral sequence correlates with the degree to which animals subsequently pursue devalued outcomes [25]. Stimulation of the DLS during behavioral initiation elicits increased habit-like behavior [26], as well as inducing reduced goal-directed control [27]. Thus, it is plausible that in the present study, the multivariate distributed predictive signal of behavioral strategy reflects neuronal populations directly involved in driving the behavioral onset of habitual actions.

Surprisingly, we did not find robust evidence for differential signatures of goal-directed and habitual control outside the posterior putamen, including in the anterior caudate or prefrontal cortex. The specificity of our findings is unexpected, given the prevailing view that additional brain regions contribute to the allocation of behavioral control across these strategies. These include areas implicated in arbitration between model-based and model-free reinforcement learning, such as the ventrolateral pre-frontal cortex and the frontopolar cortex [28], as well as areas involved in outcome monitoring and feedback-related behavioral adjustment [29, 30], such as the anterior cingulate cortex. In the present study, several reasons may explain why we did not observe decoding effects in these areas. The first is that we are focusing on a specific time window of initial learning. If those other areas are involved in active top-down control of behavioral strategy, they might not be differentially engaged during initial training, but instead become more prominently involved when the two behavioral strategies conflict markedly in their proposed action-selection policies, which would occur during the devaluation phase. This possibility could be tested, provided that trivial action-selection differences are taken into account, as noted earlier. However, in the present study, it is not feasible to reliably decode neural activity patterns during the devaluation probe because it is a brief one-shot manipulation, severely limiting the amount of data available for a decoder to utilize. To overcome these limitations, a future study could employ repeated devaluation and revaluation to yield sufficient data for robust decoding of activity associated with behavioral control during such scenarios [31]. Another possible reason for the absence of robust neural signatures of behavioral strategy in these regions is that patterns of neural activity associated with habitual and goal-directed control might not be sufficiently similar across participants to enable between-participant decoding. The posterior putamen may be unique in possessing neural coding patterns sufficiently similar across participants to allow between-participant decoding.

Literature associated the tendency to express habits behaviorally (and exhibit reduced goal-directed control) with several psychological traits and dispositions, including increased stress, anxiety, and compulsivity [14, 17, 32–34]. In the present study, because we examine behavioral performance on a single task with a one-shot devaluation procedure, we cannot ascertain whether the putamen signal we detect reflects trait-related dispositional effects of an individual’s general propensity to engage in habits and goal-directed control, or whether the putamen activity is instead indexing transient neuronal correlates of a habitual or goal-directed response tendency. Consequently, the neuronal correlates of the behavioral control strategy observed here could reflect either stable between-participant differences in neuronal encoding within the putamen or a context-specific snapshot of current task performance. However, none of the measures of individual traits, including obsessive-compulsive traits, anxiety, or mood, could explain the observed effects of devaluation sensitivity in the putamen. Thus, we can at least rule out an explanation of putamen activity in terms of the specific trait measures we obtained.

The finding that it is possible to decode an individual’s overall behavioral strategy on a task from activity measured even within a single brain area raises the possibility that brain activity patterns can potentially predict subsequent behavioral strategies and ultimately performance, in the context of brain-machine interfaces and even potentially in real-world settings. Such a capability could help assign individuals to tasks or roles for which they are best suited, providing guidance on the most appropriate learning strategy for a specific individual, or even indicating whether and when to intervene to promote the adoption of alternative learning strategies. Although the current decoding accuracy of 62% is robust and significantly above chance, it is likely to fall short of the level required for practical application. Moreover, it is infeasible to use MRI for this application due to its lack of portability and high cost. Nevertheless, the present findings provide proof of principle that activity patterns can predict even high-level behavioral strategies across individuals, opening the potential for future applications if improved methodology and less noisy, more portable brain measures can enhance out-of-sample accuracy.

Several limitations of the present study warrant consideration. First, we assessed the behavioral strategy using a single-task, one-shot outcome-devaluation procedure, which limits our ability to determine whether the decoded posterior putamen signal reflects stable, trait-like differences in behavioral control or transient, context-dependent states expressed during learning. To disambiguate these possibilities, we require longitudinal designs or tasks that incorporate repeated devaluation and revaluation to dissociate these possibilities. Second, although decoding generalized across independent healthy and psychiatric samples, the multivariate patterns identified here do not establish a causal role for posterior putamen activity in governing behavioral control. Causal inference will require interventional approaches, such as noninvasive neuromodulation or lesion studies. Third, our decoding analyses focused on activity during instrumental training. This choice avoids trivial confounds during devaluation but limits conclusions about the neural mechanisms engaged during explicit strategy conflict. Fourth, our sample was overwhelmingly right-handed (≈ 97%), precluding conclusions about whether the left posterior putamen lateralization generalizes across handedness groups. Finally, while decoding accuracy is robust and generalizable, it remains modest and below the levels required for practical application, and reliance on fMRI limits scalability due to cost and portability. These considerations highlight important directions for future work while reinforcing the present findings’ conservative and mechanistically informative account of posterior putamen involvement in behavioral control.

The present study shows that distributed activity in the human posterior putamen encodes which behavioral control strategy an individual uses during instrumental action selection. These findings provide evidence that it is possible to infer an individual’s overall behavioral control strategy from circumscribed, localized brain activity. Furthermore, the findings provide novel insights into the potential functions of the human posterior putamen, consistent with the possibility that it promotes the onset of different behavioral strategies at the time of action initiation.

## Acknowledgments

The National Institute of Mental Health supported this work under grant R01MH121089-01. We thank the Caltech Brain Imaging Center for assistance with MRI data acquisition and all participants for their involvement in the study.

## Methods

### Task Description

Participants completed the *Coin-Collector task*, a reinforcement learning task in which they used a trackball to execute leftward or rightward swipes and press buttons to collect gold and silver coins. The task comprised three sequential phases: Training, Consumption, and a Choice Test (Figure 1). Before scanning, participants received standardized, in-person instructions from a trained research assistant to ensure understanding of the task structure and response requirements. For clarity, we illustrate only the core experimental phases in the task schematic and omit the post-task contingency memory assessment.

### Training Phase

During the training phase, participants learned stimulus–action–outcome associations by performing leftward or rightward swiping actions to collect gold or silver coins. In each block, a visual cue indicated which coin was available. A brief visual feedback confirmed successful collection.

Training consisted of 20 blocks (10 per run), each lasting 37–60 seconds. Rewards were delivered according to a random-ratio schedule, promoting learning under conditions of probabilistic reinforcement, a pattern commonly associated with habit formation. Both coin types were of equal value. Actions incurred a small cost relative to successful coin collection, and we instructed the participants to collect as many coins as possible. Neural data from this training phase formed the basis for all subsequent decoding analyses.

### Consumption Phase

After the first 10 training blocks, participants completed a pre-devaluation consumption test in which they selected 10 coins from a grid containing equal numbers of gold and silver coins within 10 seconds. This phase served to familiarize participants with the consumption procedure before devaluation and to establish baseline performance, for which selecting any 10 coins was optimal. Participants then completed training blocks 11–20 using the same task structure as before.

Following training, we devalued one coin type by informing participants that the corresponding piggy bank was full. Participants subsequently completed a post-devaluation consumption test with the same coin display. We defined the *consumption score* as the number of selections directed toward the still-valued (non-devalued) coin, which provided an index of participants’ sensitivity to the devaluation manipulation.

### Choice Test Phase

In the final phase, participants completed a choice test designed to assess sensitivity to outcome devaluation. The block presented both cues simultaneously, with the associated coins hidden from view. Participants swiped freely for 90 seconds under extinction conditions (i.e., without feedback), with the instruction to maximize total points. We quantified performance using the *devaluation-ratio*, defined as the proportion of swipes directed toward the still-valued (non-devalued) outcome relative to all swipes devaluation-ratio (∈ [0, 1]), with higher values indicating greater sensitivity to the devaluation manipulation.

### Contingency Memory Assessment

Following the main task, participants completed a brief contingency knowledge test assessing explicit knowledge of task contingencies. Participants answered four binary questions: two probed stimulus–response associations (cue symbol to swiping direction), and two probed response–outcome associations (coin color to swiping direction). We didn’t provide feedback during this assessment. Furthermore, we scored responses conservatively, so a category is marked correct only if the participant answers both corresponding questions correctly. These binary measures are described as *stimulus contingency* and *coin contingency*, respectively.

### Participants

We recruited healthy adult participants (ages 18–65 years) from the Greater Los Angeles area through community-based recruitment. Participants were fluent in English, had normal or corrected-to-normal vision, met MRI safety criteria, and reported no history of neurological disorders or substance abuse. We instructed participants to abstain from recreational drugs and alcohol before scanning. All participants provided written informed consent in accordance with protocols approved by the California Institute of Technology Institutional Review Board. We compensated participants for their time at standard hourly rates for in-scanner and pre-scan procedures.

### Psychiatric Patient Sample

In addition to healthy participants, we recruited an undifferentiated sample of psychiatric patients with a diverse array of psychiatric disorders. Patients initially self-reported their diagnoses and were screened over the phone. Those who passed this screening completed an extended in-person evaluation with a clinical psychologist or psychiatrist before MRI scanning to confirm their current diagnosis. Thus all patients underwent a structured DSM-5 based structured clinical interview. In total, we recruited 68 patients spanning 12 different diagnoses (Supplementary: Table S2); diagnostic distributions in Supplementary: Table S1.

### Exclusions

Participants completed two runs of the task during fMRI scanning. We excluded eighteen participants due to incomplete imaging data. We applied the following predefined exclusion criteria to ensure interpretable behavioral performance:

1. **Low consumption score (***<* 8): indicating insufficient sensitivity to the devaluation manipulation.
2. **Devaluation ratio (***<* 0.333): reflecting responses opposite to task contingencies, rendering behavioral classification ambiguous.
3. **A low response rate during training (***<* 0.5 *swipes/s*): indicates difficulty acquiring the task.

After applying these criteria, the final sample consisted of 144 healthy controls (78 female; mean age = 30.0 ± 9.8 years) and 55 psychiatric patients (45 female; mean age = 28.2 ± 9.5 years). We present a graphical summary of participant exclusions in Supplementary: Figure S1.

### Data Analysis

We performed all data analysis in Python. We present the results as mean ± standard deviation unless reported otherwise.

### Group Classification: Beta Mixture Modeling

To classify participants based on sensitivity to outcome devaluation, we modeled the distribution of devaluation ratios using a finite mixture of Beta distributions [35], which naturally accommodate bounded data and boundary values. Models with one to five components were fit to the Initial Cohort only and evaluated using the Kolmogorov–Smirnov statistic. A two-component solution provided the most parsimonious account of the data (see Supplementary: Figure S2) and was used to define latent subgroups corresponding to low versus high sensitivity to devaluation.

We used the posterior decision boundary between components (devaluation-ratio = 0.648) to classify participants as habitual or goal-directed for all subsequent analyses. Robustness analyses indicated no evidence of systematic misfit and confirmed that downstream results were not sensitive to the specific thresholding scheme (see Supplementary: (Behavioral Classification: Beta mixture modeling of devaluation-ratio)).

### Response Rates

To assess participants’ engagement and learning during training, we analyzed response rates across task blocks. Response rate served two purposes: first, as a behavioral index of task acquisition, given that increased responding maximized reward despite participants not being explicitly instructed to do so; and second, as a potential confound, allowing us to test whether goal-directed and habitual participants differed in engagement or learning dynamics.

Response rates were analyzed using linear mixed-effects models fit separately for each training run in the Initial Cohort. Models included block number (centered within run), behavioral group (goal-directed vs. habitual), and their interaction as fixed effects, with subject modeled as a random intercept. We based the group classification on the mixture-model threshold (devaluation-ratio = 0.648). We report the fixed-effect estimates and statistical tests in the Results section. We provide the full model specification and estimation details in the Supplementary: Response Rate Analysis.

### Questionnaire Self-Reports

Participants completed multiple psychological questionnaires assessing affective, cognitive, and personality traits. Typically, we administered the questionnaires approximately one week before the fMRI session. We assessed State anxiety (STAI-State) immediately before scanning to capture transient anxiety levels. Participants who missed questionnaire sessions completed them within one week following the scan.

Psychological measures were included as confounding variables in fMRI analyses to control for individual differences in affective and cognitive state. We retained the participants with incomplete questionnaire data, and missing values in the psychological confound matrix were imputed using a K-nearest neighbors approach [36]. Although we administered the NEO Personality Inventory, we excluded it from analysis due to a coding error during data collection. We provide the complete list of questionnaires in Supplementary: Table S4.

### Logistic Regression Predicting Devaluation Sensitivity

To identify behavioral and trait-level predictors of sensitivity to outcome devaluation, we fit multivariable logistic regression models with binarized devaluation-ratio (goal-directed vs. habitual strategy) as the dependent variable. Participants were classified as goal-directed if devaluation-ratio ≥ 0.648 and habitual otherwise. Models were estimated using maximum likelihood as implemented in statsmodels (Python).

Predictors included task-related measures (response rate, task order, devalued coin identity, devalued response direction, and contingency knowledge), psychometric questionnaire scores, and demographic covariates (age, sex, and handedness). We standardized all predictors before model fitting. Missing values were imputed using a *k*-nearest neighbors approach (*k* = 5), as described above.

We assessed model fit using likelihood ratio tests and pseudo-*R*^2^ statistics. We evaluated the contribution of individual predictors using Type III Wald *χ*^2^ tests. We report the regression results in the Results section and Supplementary: Table S3.

We conducted the logistic regression analyses in two stages. First, we fit an initial model restricted to the Initial Cohort to identify behavioral and psychological variables that covaried with the devaluation-ratio and could therefore plausibly confound subsequent neural decoding analyses. This preliminary analysis informed the selection of nuisance regressors included in robustness checks of the fMRI results. Second, to more comprehensively characterize behavioral and psychometric correlates of devaluation sensitivity, we fit the same model to the full sample of healthy controls and patients. Because the full-sample analysis provides greater statistical power and broader generality, we only report these results in the Supplementary (Behavioral and Psychometric Correlates of Devaluation Sensitivity), analyses restricted to the Initial Cohort yielded qualitatively similar patterns of association with task variables.

### MRI methodology

#### Acquisition

Functional MRI data were acquired at the Caltech Brain Imaging Center (Pasadena, CA) using a Siemens Prisma 3T scanner equipped with a 32-channel radiofrequency head coil. Functional scans were collected using a multiband echo-planar imaging (EPI) sequence with 72 slices, a −30*^◦^* slice tilt relative to the AC–PC line, 192 mm × 192 mm field of view, and 2 mm isotropic resolution (TR = 1.12 s, TE = 30 ms, flip angle = 54*^◦^*, multiband acceleration = 4, in-plane acceleration factor = 2, echo spacing = 0.56 ms, EPI factor = 96).

Following each functional run, EPI-based field maps with both positive and negative phase-encoding polarities were acquired using similar parameters, with single-band acquisition (TR = 5.13 s, TE = 41.40 ms, flip angle = 90*^◦^*).

We acquired high-resolution T1-weighted and T2-weighted structural images for each participant with 0.9 mm isotropic resolution and a 230 mm × 230 mm field of view. We acquired T1-weighted images with TR = 2.55 s, TE = 1.63 ms, inversion time (TI) = 1.15 s, flip angle = 8*^◦^*, and in-plane acceleration factor = 2. We acquired T2-weighted images with TR = 3.2 s, TE = 564 ms, and in-plane acceleration factor = 2.

#### Preprocessing

We preprocessed functional MRI data using a two-stage pipeline. First, we performed anatomical and functional preprocessing using fMRIPrep (version 23.1.3), including motion correction, slice-timing correction, spatial normalization, and nuisance confound estimation.

In a second stage, we performed additional denoising and standardization using Nilearn (version 0.10.1), including a manual correction for a known z-scoring issue [37]. This stage included high-pass temporal filtering, detrending, and nuisance signal regression, including motion parameters, CompCor components, and the global signal. Preprocessed time series were standardized within run and concatenated across runs to form participant-level inputs for multivariate analyses. We provide a schematic overview of the preprocessing workflow in Supplementary: Figure S10.

#### Regions of Interest

We defined eleven regions of interest (ROIs) based on prior work implicating these regions in goal-directed and habitual control. Anatomical ROIs included bilateral anterior and posterior caudate and putamen, defined using the striatal parcellation of Pauli et al. (2018) [38].

In addition, we specified functionally defined prefrontal ROIs as 6-mm-radius spheres. A dorsolateral prefrontal cortex (dlPFC) ROI centered at (48, 9, 36) based on Gläascher et al. (2010) [39] and mirrored to the left hemisphere. A ventromedial prefrontal cortex (vmPFC) ROI centered at (−3, 41, −11) based on McNamee et al. (2013) [13]. We used all ROIs as masks for both searchlight-based and region-wise decoding analyses.

### fMRI - General Linear Modelling

We used Nilearn (version 0.10.1) [40] for all fMRI-GLM analysis.

#### First Level

We constructed first-level general linear models (GLMs) for each participant using a design matrix spanning both concatenated fMRI runs, while modeling only the training blocks. Task-related regressors were not defined for the subsequent choice-test phase, thereby preserving those time periods from contamination in downstream prediction analyses. Each run comprised 10 training blocks.

Two task-related regressors were specified: (i) a block-mean regressor capturing sustained activation within each training block, and (ii) a block-linear regressor indexing linear progression across blocks. We convolved both regressors with a canonical SPM hemodynamic response function, including temporal and dispersion derivatives [41]. The design matrix also included an intercept, yielding a total of 7 regressors (2 task regressors and their temporal and dispersion derivatives, plus the intercept). We used a 5 mm full-width-at-half-maximum Gaussian kernel to smooth functional images before analysis. Supplementary: Figure S11 provides visualizations of the design matrix and the contrasts.

#### Second Level (Parametric Analysis)

To examine how individual differences in psychological traits and task behavior related to neural activity, we performed second-level parametric analyses using participant-wise confound variables as predictors. The design matrix included standardized task-related measures, psychometric questionnaire scores, and demographic variables, with missing values imputed using a *k*-nearest neighbors approach (*k* = 5) [42, 43].

We estimated voxel-wise second-level models using mass-univariate GLMs, with each predictor of interest evaluated while controlling for all other covariates in the design matrix. Statistical contrasts isolated the effect of individual predictors. We provide a schematic overview of the second-level analysis in Supplementary: Figure S12 .

#### Statistical Inference (Activation Analysis)

For first-level statistical inference, we assessed task-related activation using voxel-wise false discovery rate (FDR) correction. Statistical maps were thresholded using FDR control applied either across the whole brain or within anatomically defined regions of interest, depending on the analysis. Results reported in the main text reflect voxels surviving these correction procedures

### Multivariate Pattern Analysis

#### Searchlight Analysis

We performed multivoxel pattern analysis (MVPA) in the Initial Cohort using a searchlight decoding approach to identify neural patterns predictive of individual differences in behavior. For each participant, first-level contrast maps corresponding to sustained training-related activity (block-mean; Supplementary: Figure S11) were used as input features. We classified participants as goal-directed or habitual based on the devaluation-ratio threshold (0.648).

We conducted searchlight decoding using a spherical searchlight (4 mm radius), with searchlight centers restricted to predefined anatomical regions of interest (ROIs) or to the whole-brain gray matter mask. We examined eleven ROIs implicated in goal-directed and habitual control. Within each searchlight, we trained a linear discriminant analysis (LDA) classifier using stratified 4-fold cross-validation, and decoding performance was quantified using classification accuracy. Furthermore, we conducted whole-brain searchlight analyses to assess whether effects extended beyond a priori regions.

#### Confound Regression

To control for nuisance effects, we performed leakage-safe confound regression within each cross-validation fold using a train-on-train residualization procedure with an intercept, following best practices for multivariate decoding [44]. We standardized confounding variables using only training-set features, and applied the estimated regression parameters on the training split to the held-out test data. This procedure ensured that no information from the test set influenced feature residualization. We provide full mathematical details of the confound regression procedure in Supplementary (Confound Regression for Multivariate Decoding).

#### Generalization Analysis

To assess out-of-sample generalization, we trained decoders on the Initial Cohort and evaluated performance in an independent held-out cohort. Unlike the discovery analysis, which used searchlight decoding, the generalization analysis used voxels within the significant decoding cluster as features.

We performed classification using an ensemble-based bagging approach with LDA as the base estimator. The ensemble comprised 100 classifiers trained on bootstrap samples, including 75% of participants and 50% of features. Final predictions were obtained by averaging predicted probabilities across ensemble members. Confound regression and feature standardization were performed using train-only statistics, as in the discovery analysis.

Because the held-out cohort was moderately class-imbalanced, we evaluated performance using the area under the receiver operating characteristic (ROC) curve (AUC), with statistical significance assessed via permutation testing of group labels.

#### Statistical Inference (MVPA)

Statistical inference for MVPA was performed on voxel-wise decoding accuracy maps using non-parametric permutation testing. A null distribution was generated by permuting class labels (5,000 permutations) and recomputing searchlight decoding maps.

We identified clusters of above-chance decoding accuracy using a cluster-forming threshold derived from the voxel-wise null distribution. Cluster-level statistics were computed by summing supra-threshold accuracy values within contiguous regions. We determined family-wise error–corrected significance by comparing observed cluster statistics to the permutation-derived null distribution of maximum cluster scores. This procedure was applied within individual ROIs and across the whole-brain gray matter mask. We provide full algorithmic details in Supplementary: Algorithm 1

#### Robustness Analyses

We conducted robustness analyses using a consistent multivariate decoding framework applied to the full dataset (*N* = 199). We trained LDA classifiers using 5-fold stratified cross-validation with repeated resampling. Within each fold, features were residualized with respect to confounds using train-only statistics and standardized before classification. Model performance was quantified using ROC AUC and averaged across folds and repetitions. This framework was applied uniformly across robustness checks to ensure comparability.

## Data and materials availability

All code used to generate the analyses and figures reported in this manuscript is publicly available at GitHub.

We provide scripts and notebooks that reproduce all figures from project-level CSV files and per-subject trial data, along with instructions for setting up the computational environment and executing the analyses. Using the provided scripts, readers can generate executed versions of all figure notebooks, ensuring full computational reproducibility. We also provide scripts to derive all task-related variables directly from the trial-level data, for analytical transparency.

We include anonymized behavioral data and derived analysis outputs (e.g., response rates, consumption scores, devaluation ratios, and decoding summaries) in the repository after removing personally identifying information. These data are sufficient to reproduce all behavioral and multivariate analyses reported in the manuscript.

Due to privacy and consent constraints, we do not publicly share raw neuroimaging data. However, we include preprocessed neuroimaging derivatives, region-of-interest masks, and analysis outputs used for multivariate decoding where permissible. We make raw neuroimaging data available to the corresponding author upon reasonable request, subject to institutional approval.

