## Supplementary for "Distributed activity in the human posterior putamen distinguishes goal-directed from habitual control"

### Supplement

#### Participant Information

This section provides additional details on participant inclusion and exclusion criteria, as well as the clinical composition. Figure S1 summarizes participant flow and exclusion criteria across healthy participants and patients. Table S1 and Table S2 report diagnostic abbreviations and the distribution of psychiatric diagnoses in the patient cohort.

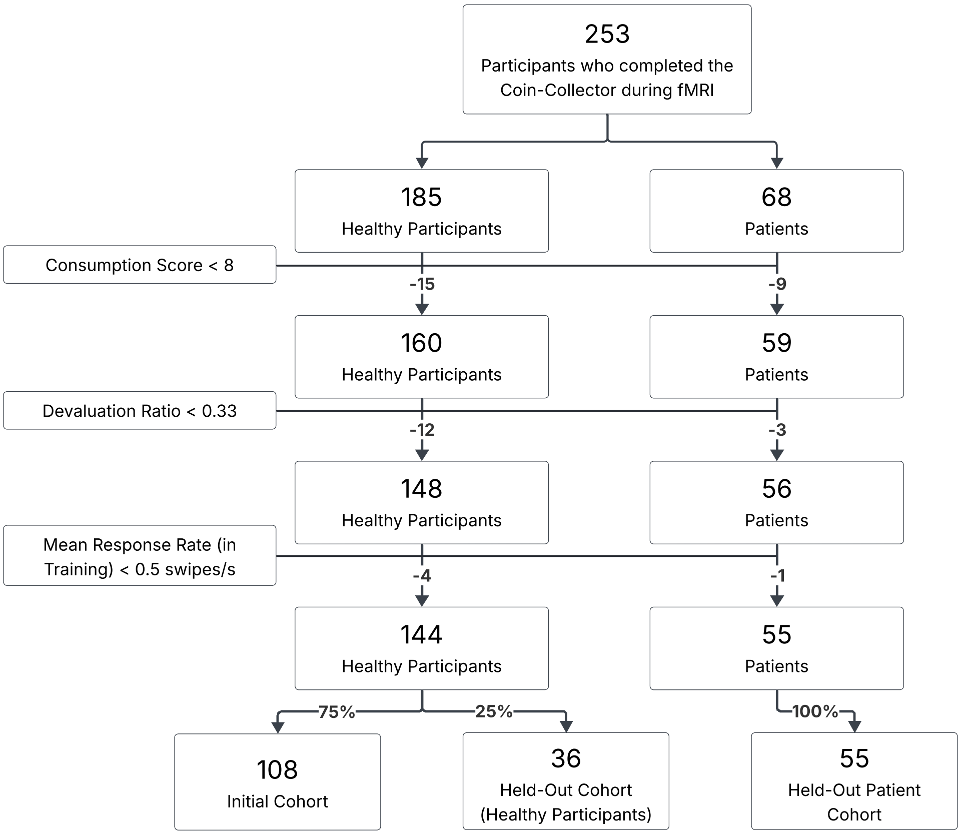


**Fig. S1**: **Participant flow and exclusion criteria.** Flow diagram showing participant inclusion and exclusion across the study. Participants were excluded sequentially based on consumption score, devaluation-ratio, and mean response rate during training. The final sample comprised an Initial Cohort, a held-out Healthy Cohort, and a held-out Patient Cohort used for generalization analyses.

**Table S1**: Clinical diagnosis abbreviations used in the study. Diagnoses were determined based on structured clinical interviews.

| Dx | Full Diagnosis Name |
|---|---|
| ADD | Attention-Deficit Disorder |
| ADHD | Attention-Deficit/Hyperactivity Disorder |
| AG | Agoraphobia |
| BDD | Body Dysmorphic Disorder |
| GAD | Generalized Anxiety Disorder |
| MDD | Major Depressive Disorder |
| MDE | Major Depressive Episode |
| OCD | Obsessive-Compulsive Disorder |
| PD | Panic Disorder |
| PMDD | Premenstrual Dysphoric Disorder |
| PTSD | Post-Traumatic Stress Disorder |
| SAD | Social Anxiety Disorder |

**Table S2**: Summary of psychiatric diagnoses among the patient sample, including total counts, proportion of participants on psychotropic medication, and proportion with at least one co-occurring diagnosis.

| Dx | Total Count | On Medication | Co-occurring Dx |
|---|---|---|---|
| ADD | 2 | 2 (100.0%) | 0 (0.0%) |
| ADHD | 8 | 6 (75.0 %) | 7 (87.5%) |
| AG | 6 | 3 (50.0 %) | 5 (83.3%) |
| BDD | 6 | 4 (66.7 %) | 5 (83.3%) |
| GAD | 35 | 19 (54.3 %) | 21 (60.0%) |
| MDD | 22 | 13 (59.1 %) | 15 (68.2%) |
| MDE | 9 | 7 (77.8 %) | 6 (66.7%) |
| OCD | 14 | 7 (50.0 %) | 9 (64.3%) |
| PD | 7 | 4 (57.1 %) | 7 (100.0%) |
| PMDD | 1 | 1 (100.0%) | 0 (0.0%) |
| PTSD | 6 | 1 (16.7 %) | 6 (100.0%) |
| SAD | 5 | 3 (60.0 %) | 4 (80.0%) |

#### Behavioral Classification: Beta mixture modeling of devaluation-ratio

To classify participants based on their sensitivity to outcome devaluation, we modeled the distribution of devaluation ratios using a finite mixture of Beta distributions. Because the devaluation-ratio is bounded in the unit interval $[0, 1]$ and includes boundary values, Beta distributions provide a natural and flexible family for capturing heterogeneity in response profiles.

***Model specification.***

We assumed that each observed devaluation-ratio $x_i$ was drawn from a convex combination of $K$ Beta distributions,

$$x_i \sim \sum_{k=1}^{K} \pi_k \, \mathrm{Beta}(\alpha_k, \beta_k), \tag{1}$$

where $\pi_k$ denotes the mixing proportion of component $k$ and $(\alpha_k, \beta_k)$ are the shape parameters governing its distribution. The corresponding mixture cumulative distribution function is

$$F_\theta(x) = \sum_{k=1}^{K} \pi_k \, F_{\mathrm{Beta}}(x \mid \alpha_k, \beta_k). \tag{2}$$

Model parameters were estimated using an Iterated Method of Moments Expectation–Maximization (IMM–EM) algorithm [35], which combines responsibility-weighted expectation steps with moment-matching maximization steps and explicitly accommodates observations at the boundaries (0 and 1). Model fitting was initialized with 1,000 random seeds using a distance-aware initialization strategy [45] to reduce sensitivity to local optima.

***Model selection.***

To determine the appropriate number of mixture components, we fit models with $K = 1$ through 5 components using data from the Initial Cohort only. We evaluated model adequacy using the Kolmogorov-Smirnov (KS) statistic,

$$D_n = \sup_{x \in [0,1]} \left| F_N(x) - F_\theta(x) \right|, \tag{3}$$

which quantifies the maximum discrepancy between the empirical cumulative distribution function $F_N(x)$ and the fitted mixture distribution. As shown in Figure S2 (A), model fit improved substantially from one to two components and plateaued thereafter. We therefore selected a two-component solution as the most parsimonious representation of the data.

***Parameter estimates and classification threshold.***

The two-component model identified one component centered near intermediate devaluation ratios and another near high devaluation ratios, corresponding to habitual and goal-directed response profiles, respectively (Figure S2). The estimated parameters were $(\alpha, \beta) = (3.375, 0.460)$ for the goal-directed component and $(\alpha, \beta) = (29.636, 28.435)$ for the habitual component, with mixing proportions $\pi = 0.541$ and $\pi = 0.459$, respectively. We used the posterior decision boundary separating the two components, located at devaluation-ratio $= 0.648$, to classify participants in all subsequent analyses.

***Generalization across cohorts.***

To assess generalizability, we overlaid the mixture model fitted to the Initial Cohort onto DR distributions from independent cohorts, without refitting the model's parameters. As shown in Figure S2 (C-D)), the bimodal structure of devaluation-ratio values was preserved in both a held-out healthy cohort and a heterogeneous patient cohort, indicating that the two-component solution captures a stable behavioral phenotype rather than cohort-specific noise.

***Goodness-of-fit and robustness.***

We further evaluated model robustness using a cross-validated parametric bootstrap procedure based on the KS statistic (5,000 replicates). For each replicate, we generated synthetic datasets from the fitted model, re-estimated the parameters on bootstrap training samples, and computed KS distances on held-out test samples. In the Initial Cohort, the mean KS distance was 0.1639, with a bootstrap $p$-value of 0.41, indicating no evidence of systematic misfit. Comparable results were obtained in the full healthy control sample ($n = 144$; mean KS distance $= 0.1544$, bootstrap $p = 0.28$), supporting the robustness of the two-component solution.

Finally, we confirmed that downstream decoding results were not sensitive to the specific binarization strategy by comparing mixture-based classification with alternative thresholding schemes (median split and quartile-based contrasts; Table S3)), which yielded comparable decoding performance.

#### Response Rate Analysis

This analysis assessed learning-related changes in response vigor during training and evaluated whether differences in response rate could confound subsequent group comparisons and neural decoding analyses.

***Model specification.***

Response rates were analyzed using linear mixed-effects models fit separately for each training run in the Initial Cohort. We classified participants as goal-directed or habitual using the mixture-model threshold derived from the devaluation-ratio (0.648).

For participant $s$, block $i$, and run $r$, response rate was modeled as:

$$\text{rate}_{isr} = \beta_0 + \beta_1\,\text{blockid}_{isr} + \beta_2\,\mathbb{I}[g_s = \text{HB}] + \beta_3\,\text{blockid}_{isr} \cdot \mathbb{I}[g_s = \text{HB}] + b_{0s} + \varepsilon_{isr}, \quad (4)$$

where we treated block number as a continuous predictor (centered within run), we included group as a categorical fixed effect (goal-directed as reference), and $b_{0s}$ denotes a subject-level random intercept. Random intercepts were assumed to follow a normal distribution, $b_{0s} \sim \mathcal{N}(0, \sigma_b^2)$, and residual errors were assumed to be normally distributed, $\varepsilon_{isr} \sim \mathcal{N}(0, \sigma^2)$.

***Estimation and inference.***

Models were fit using restricted maximum likelihood (REML) as implemented in the `statsmodels` Python package, with optimization performed via the L-BFGS

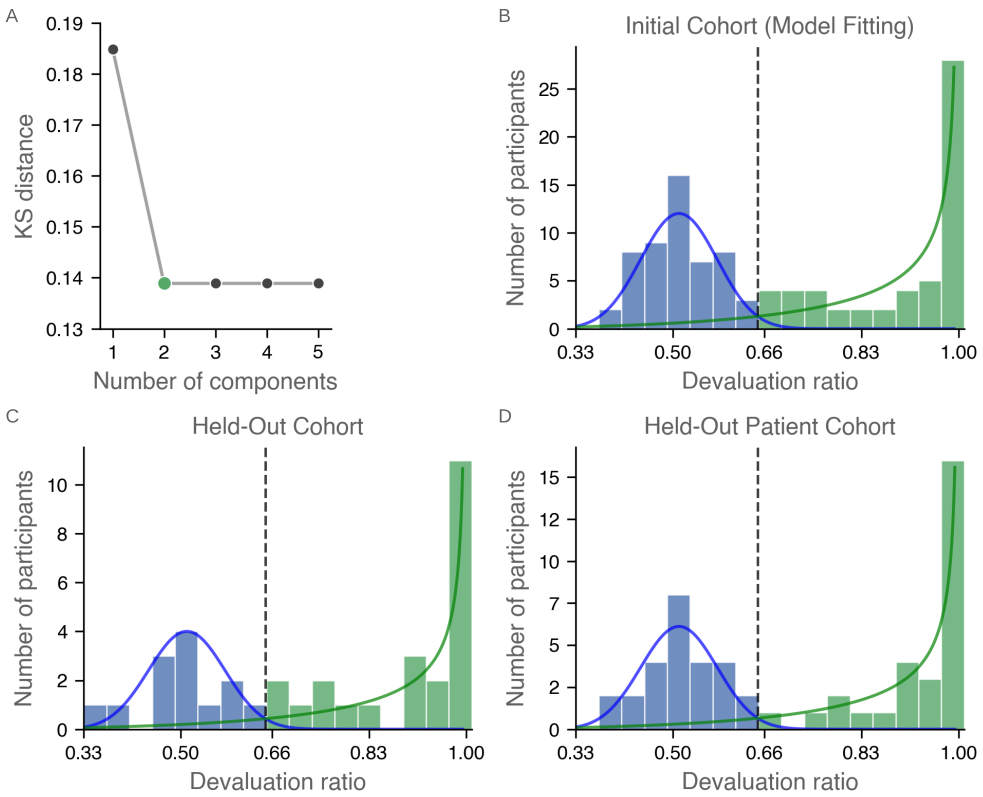


**Fig. S2**: **Beta mixture modeling of devaluation-ratio. (A)** Kolmogorov–Smirnov (KS) distance as a function of the number of Beta-mixture components fit to the Initial Cohort, showing a marked improvement from one to two components and a plateau thereafter. **(B)** Distribution of devaluation-ratio values in the Initial Cohort with the fitted two-component Beta mixture overlaid; curves show the weighted mixture components and their sum, scaled to expected participant counts. The dashed vertical line indicates the mixture-derived decision boundary (threshold = 0.648). **(C)** Devaluation ratio distribution in a held-out healthy cohort with the Initial-Cohort mixture model overlaid. **(D)** Devaluation ratio distribution in a held-out patient cohort with the same model overlaid, demonstrating preservation of the bimodal structure across populations.

algorithm (maximum 1000 iterations). We conducted statistical inference using fixed-effects Wald $z$-tests. We derived group-specific learning slopes by combining the main effect of block number with the group $\times$ block interaction term.

***Variance components.***

The fitted models revealed reliable between-subject variability in baseline response rates in both training runs, indexed by non-zero random intercept variance (Run 1: $\sigma_b^2 = 0.771$; Run 2: $\sigma_b^2 = 0.983$).

***Relation to main analyses.***

This response-rate analysis served as a control to characterize learning dynamics and participant engagement during training. Mean response rate and learning slopes (for each run) were subsequently included as nuisance regressors in multivariate decoding analyses to ensure that differences in motor output or response vigor did not drive neural classification results.

#### Behavioral and Psychometric Correlates of Devaluation Sensitivity

To characterize behavioral and trait-level correlates of individual differences in devaluation sensitivity, we fit a multivariable logistic regression model predicting binarized devaluation-ratio (goal-directed vs. habitual strategy) from task-related measures, psychometric questionnaire scores (Table S4), and demographic variables (Figure S3, Table S3). We standardized all predictors before analysis, and missing values were imputed using a $k$-nearest neighbors approach ($k = 5$).

Among task-related variables, coin-contingency sensitivity emerged as the strongest predictor of goal-directed behavior, showing a significant positive association with devaluation sensitivity. Response rate showed a trend-level positive association in the same direction, whereas other task measures did not contribute significantly.

Within the psychometric domain, obsessive-compulsive symptom severity (OCI-R total score) was the only measure that significantly predicted behavior, with higher scores associated with a greater likelihood of habitual responding. No other psychometric scale showed a reliable association with devaluation sensitivity after accounting for task variables. Demographic factors, including age, sex, and handedness, were not significantly related to strategy use.

For visualization purposes, we omit psychometric predictors with minimal effects ($p > 0.25$) from Figure S3, though all predictors were retained in the statistical model and reported in full in Table S3. Together, these results indicate that individual differences in devaluation sensitivity are primarily explained by task-specific contingency knowledge, with a more limited contribution from trait compulsivity.

#### Generalization to a Held-Out Patient Cohort

To assess whether the posterior putamen decoding effect generalizes beyond healthy participants, we evaluated classifier performance in an independent cohort of psychiatric patients, held out from all stages of model development. This Held-Out Patient Cohort comprised 55 participants with heterogeneous psychiatric diagnoses (Table S2) who completed the same task under identical acquisition parameters.

We classified participants as goal-directed or habitual using the same devaluation-ratio threshold derived exclusively from the Initial Cohort. This procedure yielded

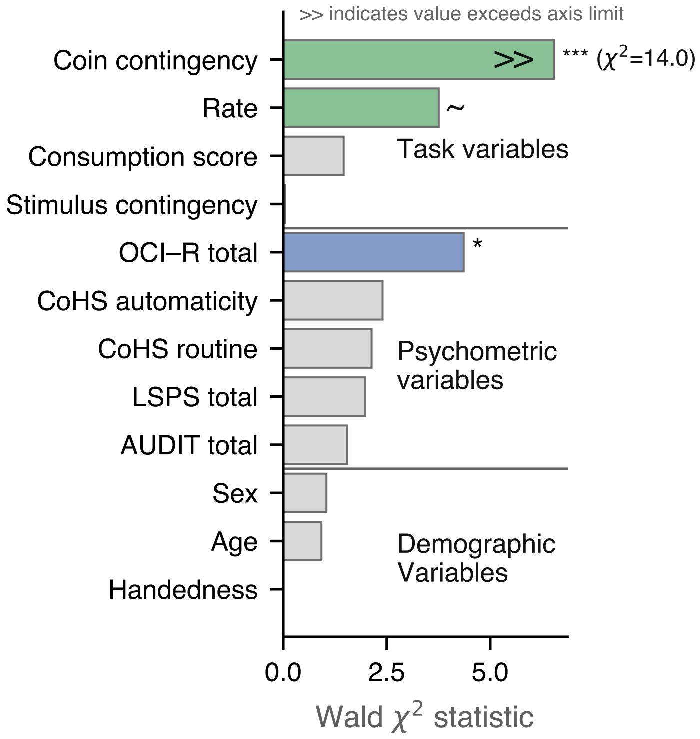


**Fig. S3**: **Behavioral, psychometric, and demographic predictors of goal-directed vs. habitual strategy.** Wald $\chi^2$ statistics from a logistic regression predicting binarized devaluation sensitivity (goal-directed vs. habitual; $N = 199$). Predictors included task-related measures, psychometric questionnaire totals, and demographic variables; we standardized all predictors, and imputed missing values using $k$-nearest neighbors ($k = 5$). Bars are ordered within domains by effect size. Green bars indicate predictors associated with greater goal-directed behavior, whereas blue bars indicate predictors associated with greater habitual behavior; gray bars indicate non-significant predictors. In the figure, we omit psychometric variables with minimal effects ($p > 0.25$) for visual clarity. We truncate the x-axis to improve readability; the double arrow (>>) denotes values exceeding the axis limit, with the corresponding $\chi^2$ value reported explicitly. Significance markers denote $p < 0.001$ (***), $p < 0.05$ (*), and $p < 0.10$ ($\sim$).

26 goal-directed and 29 habitual participants (Figure S4A). No parameters of the behavioral classification model were refit using patient data.

We applied a region-based decoder restricted to the left posterior putamen significant cluster identified in the Initial Cohort. As in the healthy held-out analysis, the decoder was trained exclusively on the Initial Cohort and applied directly to the

**Table S3**: **Logistic regression predicting goal-directed versus habitual strategy.** Logistic regression results predicting binarized devaluation sensitivity (goal-directed vs. habitual; $n = 199$). We standardize predictors before analysis. Wald $\chi^2$ statistics correspond to Type III tests with 1 degree of freedom.

| **Domain** | **Predictor** | $\beta$ | **SE** | $\chi^2$ | $p$ |
|---|---|---|---|---|---|
| Task | Consumption score | 0.194 | 0.161 | 1.46 | 0.227 |
| | Rate | 0.335 | 0.173 | 3.76 | 0.053 |
| | Coin contingency | 0.713 | 0.191 | 13.99 | $< .001$ |
| | Stimulus contingency | -0.036 | 0.168 | 0.05 | 0.831 |
| Psychometric | OCI–R total | -0.468 | 0.224 | 4.36 | 0.037 |
| | CoHS routine | -0.281 | 0.193 | 2.13 | 0.144 |
| | CoHS automaticity | 0.277 | 0.179 | 2.40 | 0.121 |
| | SRS total | 0.247 | 0.343 | 0.52 | 0.471 |
| | OLIFE total | 0.198 | 0.303 | 0.43 | 0.513 |
| | STAI total | -0.205 | 0.389 | 0.28 | 0.598 |
| | LSPS total | -0.355 | 0.253 | 1.97 | 0.160 |
| | AUDIT total | 0.207 | 0.167 | 1.54 | 0.215 |
| | EAT total | -0.111 | 0.181 | 0.38 | 0.539 |
| | PDSS total | 0.159 | 0.202 | 0.62 | 0.431 |
| | PSWQ total | 0.199 | 0.318 | 0.39 | 0.531 |
| | BDI total | -0.121 | 0.292 | 0.17 | 0.678 |
| | Apathy total | 0.084 | 0.248 | 0.11 | 0.736 |
| | BIS/BAS BIS | 0.228 | 0.278 | 0.67 | 0.412 |
| Demographic | Age | 0.168 | 0.175 | 0.92 | 0.337 |
| | Sex | 0.190 | 0.186 | 1.04 | 0.307 |
| | Handedness | -0.004 | 0.163 | 0.00 | 0.979 |

patient cohort without retraining or parameter tuning. We used a bagging ensemble of linear discriminant classifiers to reduce variance and enhance generalization.

Decoder performance in the patient cohort was quantified using ROC AUC. The decoder achieved robust above-chance performance (mean ROC AUC $= 0.739 \pm 0.019$ SD), significantly exceeding the permutation-based null distribution generated from 5,000 label shuffles ($p = 0.001$; Figure S4B). This result demonstrates that the multivariate activity pattern within the posterior putamen that distinguishes goal-directed from habitual behavior generalizes to a diagnostically heterogeneous patient population.

#### Robustness and Reproducibility of Multivariate Decoding

To assess the robustness of our multivariate decoding results, we performed a series of complementary control analyses designed to rule out methodological artifacts, overfitting, or dependence on specific analytic choices. These analyses examined the stability of decoding performance across alternative pipelines, classifier choices, group-definition schemes, cross-validation strategies, and potential task-administration confounds. Unless otherwise noted, all robustness analyses used all participants (n =

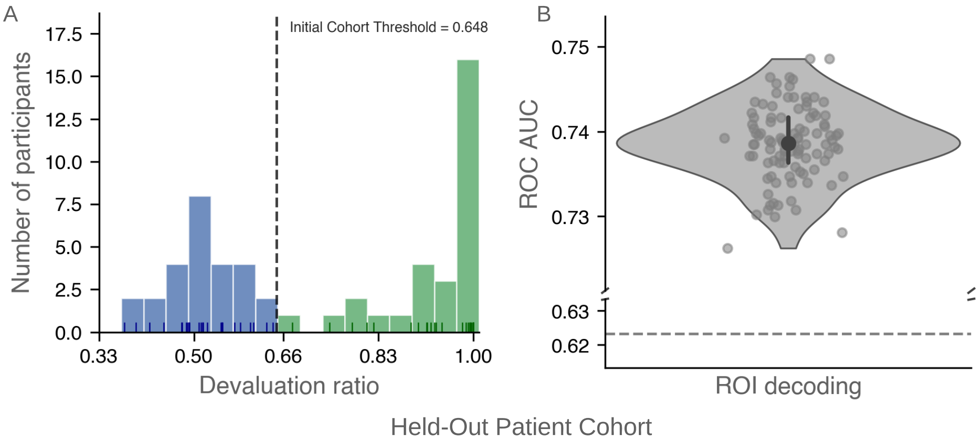


**Fig. S4**: **Generalization of posterior putamen decoding to a held-out patient cohort.** **(A)** Distribution of devaluation ratios in the held-out patient cohort (n = 55), classified using the same devaluation-ratio threshold (DR = 0.648) derived from the Initial Cohort. We classified participants with devaluation ratios above the threshold as goal-directed and those below as habitual. **(B)** ROI-level decoding performance in the held-out patient cohort obtained by applying the decoder trained exclusively on the Initial Cohort to the patient data without retraining. The violin plot shows the distribution of ROC AUC values across cross-validation repetitions. The central point denotes the mean ROC AUC, and the vertical bar indicates the interquartile range (25th–75th percentile). The dashed horizontal line indicates chance performance based on the permutation-derived null distribution. Decoding performance was significantly above chance (p = 0.001).

199), identical preprocessing, confound regression with rate and rate slopes (individually for both runs), and cross-validation procedures as in the primary ROI decoding analysis.

***Conventional MVPA pipeline.***

As a baseline comparison, we repeated the decoding analysis using a conventional MVPA pipeline commonly employed in the literature, without an explicitly held-out generalization cohort. Using cross-validated decoding with permutation-based voxel-wise inference and small-volume family-wise error rate correction within an anatomical left posterior putamen ROI, we again observed significant decoding of goal-directed vs. habitual strategy in both the whole healthy cohort and the combined sample including patients (Figure S5).

Importantly, the spatial localization of this decoding effect closely matched that identified using our primary generalization-focused pipeline. To quantify spatial reproducibility, we compared decoding clusters across three samples—the Initial Cohort ($n$ = 108), Healthy Cohort ($n$ = 144), and Full Sample ($n$ = 199)—and observed substantial voxel-wise overlap. Pairwise Dice coefficients ranged from 0.56 to 0.71, with corresponding Jaccard indices between 0.39 and 0.55, and a three-way overlap of 39 voxels across all samples. These results demonstrate that even under a stereotypical MVPA approach, the posterior putamen decoding effect is statistically robust and spatially reproducible across independent cohort definitions.

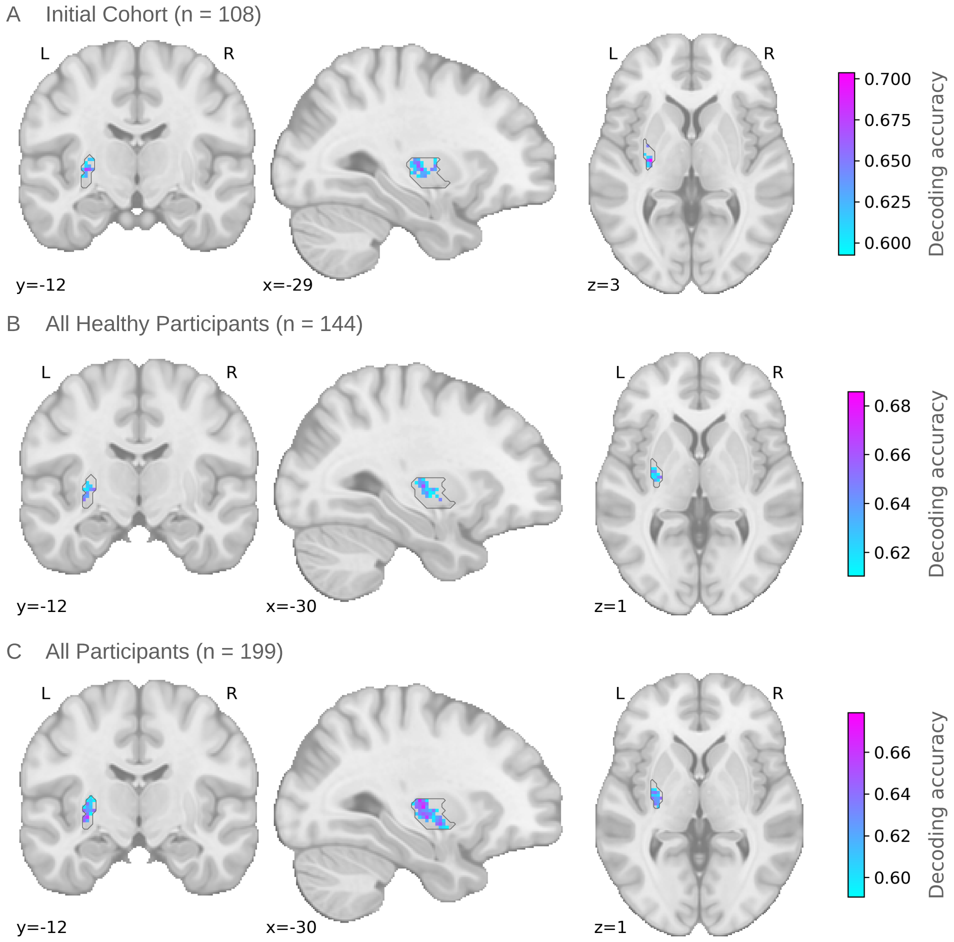


**Fig. S5**: **Reproducibility of posterior putamen decoding across cohorts.** Decoding accuracy maps for goal-directed vs. habitual strategy in the left posterior putamen across three increasingly inclusive samples: **(A)** Initial Cohort ($n = 108$), **(B)** All Healthy Participants ($n = 144$), and **(C)** All Participants, including patients ($n = 199$). Maps show cross-validated multivariate decoding accuracy within the posterior putamen, overlaid on the MNI template brain. All panels use a standard color scale to facilitate direct comparison across cohorts. Consistent spatial localization of decoding effects across samples demonstrates robustness and reproducibility of the posterior putamen signal, independent of cohort definition.

**Table S4: Decoder performance across classifier implementations in held-out cohorts.** Mean ROC AUC ± SD for decoding goal-directed versus habitual strategy within the left posterior putamen. We train all models on the Initial Cohort and evaluate them on the Held-Out Cohort (healthy participants) and the Held-Out Patient Cohort, without retraining. Bagging ensembles were used in the primary analyses (in bold) to enhance generalization, but are shown here alongside standalone classifiers to demonstrate that decoding performance does not depend on a specific classifier choice.

| Classifier | Ensemble | Healthy | Patients |
|---|---|---|---|
| Logistic regression (L1) | Standalone | 0.614 ± 0.050 | 0.628 ± 0.046 |
| Logistic regression (L1) | Bagged | 0.657 ± 0.033 | 0.688 ± 0.034 |
| Logistic regression (L2) | Standalone | 0.626 ± 0.054 | 0.634 ± 0.043 |
| Logistic regression (L2) | Bagged | 0.654 ± 0.035 | 0.682 ± 0.036 |
| Linear discriminant analysis | Standalone | 0.646 ± 0.020 | 0.734 ± 0.017 |
| **Linear discriminant analysis** | **Bagged** | **0.666 ± 0.020** | **0.739 ± 0.019** |
| Gaussian Naive Bayes | Standalone | 0.685 ± 0.024 | 0.676 ± 0.033 |
| Gaussian Naive Bayes | Bagged | 0.710 ± 0.023 | 0.651 ± 0.034 |

##### *Anatomical specificity across decoding approaches*

To assess the anatomical specificity and robustness of the decoding effect, we evaluated classification performance using both searchlight-based MVPA and region-wise decoding analyses. In the searchlight analysis, conducted across the whole brain and within predefined striatal and prefrontal regions of interest, only the left posterior putamen exhibited a cluster that survived permutation-based cluster-level correction. No other cortical or subcortical region showed reliable above-chance decoding.

To further test whether this effect depended on local searchlight parameters or spatial scale, we repeated the analysis using whole-region decoding within anatomically defined ROIs. Consistent with the searchlight results, only the left posterior putamen ROI showed significant decoding of goal-directed versus habitual strategy. In contrast, all other striatal and prefrontal regions remained at chance levels (Figure S6). Together, these converging analyses demonstrate that decoding performance is both anatomically specific and robust across multivariate analysis approaches.

##### *Robustness to Alternative Performance Metrics*

To evaluate whether decoding performance depended on the choice of evaluation metric, we re-scored decoder outputs using multiple complementary classification metrics, including ROC AUC, accuracy, balanced accuracy, precision, recall, F1 score, average precision, and Brier score loss (Table S5). We compute all metrics using identical cross-validation folds, classifier parameters, and feature sets as in the primary analyses, ensuring that differences across metrics reflect only the scoring function rather than changes in the underlying model or data.

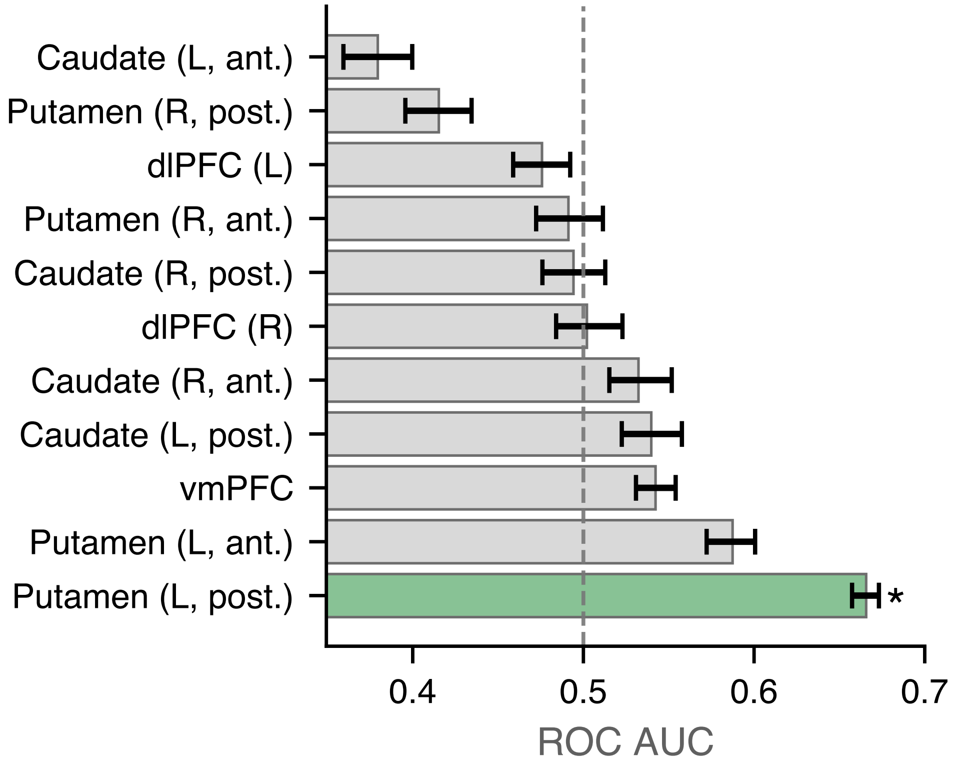


**Fig. S6**: **ROI-based decoding across anatomically defined regions.** Mean ROC AUC for decoding goal-directed versus habitual strategy across anatomically defined regions of interest. Error bars indicate the interquartile range (25th–75th percentile) across 100 repetitions of cross-validation. The dashed vertical line denotes chance-level performance (ROC AUC = 0.5). Only the left posterior putamen (green) showed reliable above-chance decoding ($p < 0.05$ (*)). In contrast, all other regions remained near chance (gray), demonstrating that decoding performance is anatomically specific and not driven by searchlight configuration.

Across metrics, decoding performance was consistently above chance and exhibited a stable central tendency. Discrimination-based metrics yielded closely aligned estimates of performance: ROC AUC and balanced accuracy both showed a mean value of 0.642 (SD $\approx$ 0.014), with 25th–75th percentile ranges spanning approximately 0.63–0.65. Overall accuracy was similarly elevated (mean = 0.645, SD $\approx$ 0.014). Metrics emphasizing positive-class performance showed comparable results, including precision (mean = 0.669, SD $\approx$ 0.013), recall (mean = 0.679, SD $\approx$ 0.019), and F1 score (mean = 0.671, SD $\approx$ 0.015). Average precision, which summarizes precision–recall trade-offs across thresholds, likewise exceeded chance (mean = 0.628, SD $\approx$ 0.011).

**Table S5**: **Robustness of posterior putamen decoding across performance metrics.** We evaluated decoder performance using multiple classification metrics. Values reflect mean performance across cross-validation repetitions, with standard deviation (SD) and interquartile range (25th–75th percentile) reported. We compute all metrics using identical data splits, feature sets, and classifier parameters to those used in the primary analyses (in bold).

| Metric | Mean | SD | 25th | 75th |
|---|---|---|---|---|
| **ROC AUC** | **0.642** | **0.014** | **0.630** | **0.652** |
| Balanced accuracy | 0.642 | 0.014 | 0.630 | 0.652 |
| Accuracy | 0.645 | 0.014 | 0.633 | 0.653 |
| Precision | 0.669 | 0.013 | 0.660 | 0.678 |
| Recall | 0.679 | 0.019 | 0.664 | 0.691 |
| F1 score | 0.671 | 0.015 | 0.660 | 0.681 |
| Average precision | 0.628 | 0.011 | 0.620 | 0.635 |
| Brier score loss | 0.355 | 0.014 | 0.347 | 0.367 |

Calibration-based assessment using Brier score loss also indicated reliable performance, with values substantially below the chance-level baseline (mean = 0.355, SD $\approx$ 0.014), consistent with non-random probabilistic predictions.

Importantly, although absolute values differed across metrics due to differences in scaling and sensitivity to class prevalence, the qualitative pattern of results was unchanged: posterior putamen activity reliably distinguished goal-directed from habitual participants across all evaluated metrics. These findings indicate that idiosyncrasies of a particular performance metric do not drive the decoding effect, but instead reflect stable separability in the underlying multivariate neural signal.

###### *Robustness to Cross-Validation Scheme*

To assess whether decoding performance depended on the specific cross-validation scheme, we repeated the ROI-level decoding analysis using a range of $k$-fold cross-validation splits ($k$ = 3–10). All analyses used identical data, feature sets, classifiers, and permutation procedures, differing only in the number of folds (Table S6).

Across cross-validation schemes, decoding performance remained stable and consistently above chance. Mean ROC AUC values ranged from 0.657 to 0.681, with overlapping interquartile ranges across all values of $k$. No systematic trend in performance was observed as a function of the number of folds, indicating that the decoding effect is not sensitive to the particular cross-validation partitioning strategy.

##### Robustness to Group-Definition Thresholds

To assess whether decoding performance depended on the specific behavioral group-definition threshold, we repeated the posterior putamen decoding analysis using alternative classification schemes. We observed comparable performance across a

**Table S6**: **Robustness of posterior putamen decoding across cross-validation schemes.** Decoder performance evaluated using different $k$-fold cross-validation splits. Values reflect mean ROC AUC across repetitions, with standard deviation (SD) and interquartile range (25th–75th percentile) reported. All analyses used identical data, classifiers, and preprocessing pipelines, differing only in the number of folds.

| CV folds ($k$) | Mean | SD | 25th | 75th |
|---|---|---|---|---|
| 3 | 0.657 | 0.024 | 0.638 | 0.674 |
| 4 | 0.669 | 0.021 | 0.658 | 0.685 |
| 5 | 0.668 | 0.016 | 0.658 | 0.678 |
| 6 | 0.672 | 0.016 | 0.663 | 0.680 |
| 7 | 0.674 | 0.015 | 0.664 | 0.683 |
| 8 | 0.676 | 0.015 | 0.666 | 0.685 |
| 9 | 0.680 | 0.016 | 0.672 | 0.688 |
| 10 | 0.681 | 0.012 | 0.673 | 0.690 |

mixture-model–derived cutoff, a median split, and a top-versus-bottom quartile split, despite differences in sample size and class balance (Table S7). These results indicate that the decoding effect is robust to reasonable variations in how goal-directed and habitual groups are defined and does not depend on a particular thresholding choice.

**Table S7**: Robustness of posterior putamen decoding across alternative group-definition schemes.

| Type of Analysis | Total $N$ | $N_{\text{Goal-Directed}}$ | Mean ROC AUC | SD |
|---|---|---|---|---|
| Mixture threshold (cutoff = 0.648) | 199 | 107 | 0.676 | 0.010 |
| Median split | 199 | 100 | 0.679 | 0.009 |
| Quartile split (top vs. bottom) | 100 | 52 | 0.648 | 0.020 |

#### Graceful Degradation of Decoder Performance

To further assess the robustness of the posterior putamen decoder, we examined how decoding performance changes as we progressively reduce the amount of training data and spatial information. Rather than collapsing abruptly, performance degraded smoothly across multiple forms of data reduction, consistent with a distributed and stable multivariate signal.

First, we evaluated performance as a function of the proportion of subjects used for training. As the subject ratio decreased, classification performance declined gradually, with no evidence of catastrophic failure even at low training fractions (Figure S7 (A)). Accuracy was reported for this analysis because, at minimal subject ratios, some cross-validation folds contained only a single class, rendering ROC AUC undefined.

It reflects a known limitation of ROC-based metrics in extremely undersampled or imbalanced settings rather than instability of the decoder itself.

Second, we assessed sensitivity to voxel subsampling by training the decoder on progressively smaller fractions of voxels within the left posterior putamen ROI. ROC AUC decreased smoothly as we reduced the voxel ratio, indicating that decoding performance does not depend on a small set of highly specific voxels but instead reflects information distributed across the region (Figure S7 (B)).

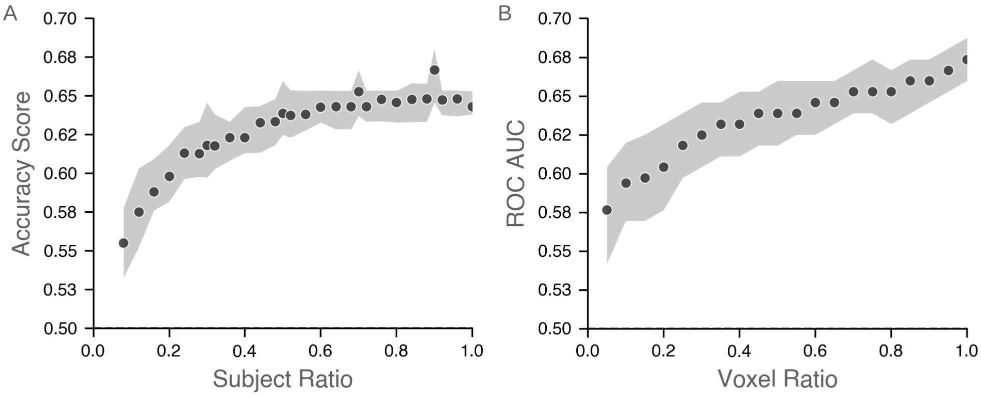


**Fig. S7**: **Graceful degradation of decoder performance with reduced training data. (A)** Classification accuracy as a function of the proportion of subjects used for training. **(B)** ROC AUC as a function of the proportion of voxels included in the analysis. Points denote median decoding performance across repetitions at each ratio, and shaded regions indicate the interquartile range (25th–75th percentile). In both panels, performance decreases smoothly as we reduce the training data, rather than collapsing abruptly, consistent with a distributed, robust multivariate signal. Accuracy is reported in Panel A because, at minimal subject ratios, some cross-validation folds contained only a single class, rendering ROC AUC undefined.

We next examined the effect of spatial smoothing on decoding performance. Increasing the Gaussian smoothing kernel led to a monotonic reduction in ROC AUC (Figure S8 (A)), consistent with the loss of fine-grained multivariate structure under spatial blurring. Similarly, systematically knocking out disjoint sets of voxels ranked by decoder weight signal-to-noise ratio resulted in a graded decline in performance (Figure S8 (B)), further supporting the contribution of distributed voxel-level information.

Finally, we assessed the stability of voxel weights across cross-validation repetitions. The mean classifier weights for individual voxels showed little variability. They maintained a consistent sign across repetitions (Figure S9), indicating that the decoder relies on a stable multivariate pattern rather than on noise-driven or fold-specific fluctuations.

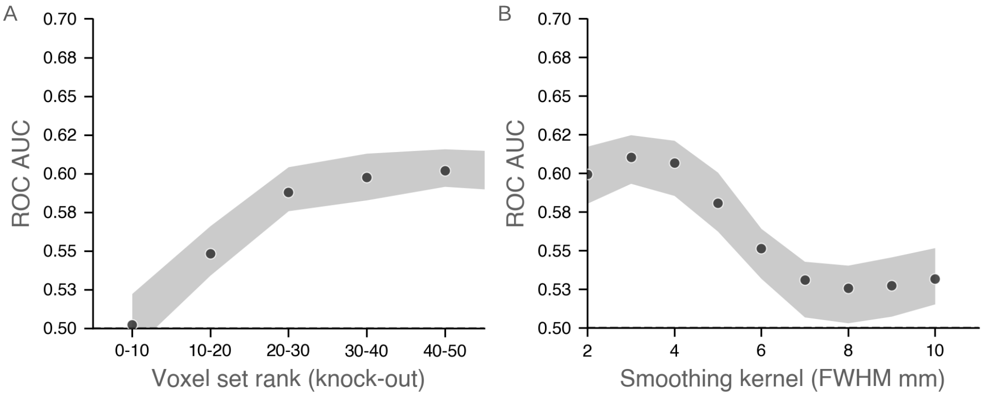


**Fig. S8**: **Decoding depends on fine-grained multivariate information. (A)** ROC AUC as a function of progressively removing disjoint voxel sets ranked by decoder-weight importance. Points denote median decoding performance across repetitions, and shaded regions indicate the interquartile range (25th–75th percentile). Removing increasingly informative voxel sets leads to graded reductions in performance, consistent with a distributed multivariate representation. **(B)** ROC AUC as a function of spatial smoothing applied to the fMRI data (Gaussian kernel, full width at half maximum). Increasing spatial smoothing leads to a monotonic decrease in decoding performance, indicating that fine-grained multivariate structure is progressively lost as spatial detail is blurred. For both analyses, the univariate signal was removed before decoding, ensuring that observed effects reflect changes in multivariate information rather than residual univariate activity.

Together, these analyses demonstrate that posterior putamen decoding performance degrades gracefully with reductions in training data and spatial information, providing convergent evidence that the observed decoding effect reflects a robust, distributed multivariate neural representation.

#### Psychometric Questionnaires

Table S8 lists all psychometric questionnaires included as covariates in the behavioral and neuroimaging analyses, along with their full names and primary references. Questionnaire scores were used as standardized predictors and nuisance covariates, as described in the Methods section.

#### Technical Issues During Task Administration: Details and Validation

##### *Description of Technical Issues*

Two minor technical issues affected response registration in subsets of participants:

- **Issue 1:** Trackball Position Reset (affected n = 25 participants after exclusions). During the choice phase, the trackball position resets after each recorded response.

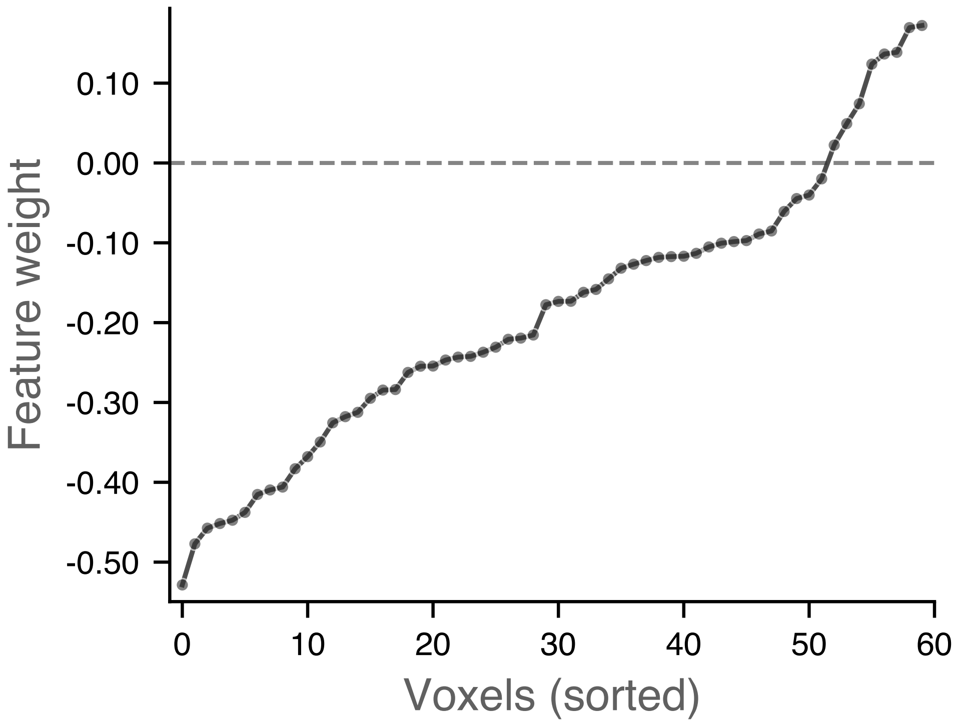


**Fig. S9**: **Stability of voxel weights across cross-validation repetitions.** Median classifier weights for the 60 voxels within the left posterior putamen cluster, averaged across cross-validation folds and sorted by median weight. Points indicate the median weight across repetitions, and shaded regions denote the full range (minimum to maximum) of voxel weights observed across cross-validation repetitions. The dashed horizontal line represents zero weight. The preserved ordering and consistent sign of voxel weights across repetitions indicate that decoding relies on a stable voxel-level pattern rather than fold-specific or noise-driven fluctuations.

If the trackball retained sufficient momentum after a swipe, this reset could trigger an unintended response in the same direction. Critically, this issue was specific to the choice phase and did not affect training-phase data used in fMRI analyses.

- **Issue 2:** Mouse Sensitivity Setting (affected n = 52 participants after exclusions). The stimulus presentation computer operated at a faster mouse sensitivity setting than intended for a subset of participants due to an inadvertent configuration error.

The increased sensitivity led it to register spurious trailing responses as multiple swipes during both the training and choice phases.

***Preprocessing Corrections***

To address these issues, we applied the following response-cleaning procedures:

**Table S8: Psychometric questionnaires included in the study.**

| Abbreviation | Full name | Reference |
|---|---|---|
| OCI–R | Obsessive–Compulsive Inventory–Revised | [46] |
| CoHS | Creature of Habit Scale | [47] |
| SRS | Self–Regulation Scale | [48] |
| OLIFE | Oxford–Liverpool Inventory of Feelings and Experiences | [49] |
| STAI | State–Trait Anxiety Inventory | [50] |
| LSPS | Liebowitz Social Phobia Scale | [51] |
| AUDIT | Alcohol Use Disorders Identification Test | [52] |
| EAT | Eating Attitudes Test | [53] |
| PDSS | Panic Disorder Severity Scale | [54] |
| PSWQ | Penn State Worry Questionnaire | [55] |
| BDI | Beck Depression Inventory | [56] |
| Apathy | Apathy Evaluation Scale | [57] |
| BIS/BAS | Behavioral Inhibition / Activation Scales | [58] |

- **Issue 1 (choice phase only):** We removed trailing responses with inter-response intervals $< 0.084$ s when the preceding response was in the same direction.
- **Issue 2 (training and choice phases):** We removed trailing responses with inter-response intervals $< 0.034$ s under the same directional criterion.

We selected these thresholds to eliminate physically implausible response sequences while preserving legitimate rapid responses.

*Impact Assessment*

**Behavioral Measures**: We tested whether key behavioral measures differed between affected and unaffected participants for Issue 2 (which impacted both training and choice phases):

- **Response rates during training:** No significant difference between participants with versus without fast mouse sensitivity ($t = 1.24$, $p = 0.22$).
- **Devaluation sensitivity:** No significant difference in devaluation-ratio between groups ($t = -0.95$, $p = 0.35$).

For Issue 1, the trackball reset affected both swiping directions equally during the choice phase, meaning the devaluation-ratio, our primary behavioral measure, would be unbiased by this artifact even before correction.

**fMRI Decoding Results**: To verify that neural decoding results, we performed two validation analyses:

- **Confound regression approach:** We included separate binary indicator variables for both technical issues in the confound regression procedure. Decoding performance remained robust (ROC AUC = 0.69), indicating that the neural signature of behavioral strategy was independent of these technical artifacts.
- **Cross-generalization test:** We trained the decoder exclusively on unaffected participants ($n = 122$) and tested on affected participants ($n = 77$). The decoder generalized successfully (ROC AUC $= 0.65 \pm 0.02$ SD), demonstrating that the

identified neural patterns were not specific to the unaffected subsample and were robust to the presence of these technical issues.

***Conclusion***

We corrected both technical issues through preprocessing. Despite Issue 2 affecting training-phase data used in fMRI analyses, validation analyses confirmed that neither issue impacted our primary behavioral or neural findings. We retain all affected participants in the final analyses.

#### Confound Regression for Multivariate Decoding

To remove nuisance variance from voxel-wise features while avoiding information leakage, confound regression was performed separately within each cross-validation fold using a train-on-train residualization procedure with an intercept [44].

Let $\mathbf{X} \in \mathbb{R}^{N\times v}$ denote the matrix of voxel-wise features across $N$ participants and $v$ voxels, and let $\mathbf{C} \in \mathbb{R}^{N\times k}$ denote the corresponding confound matrix. For a given cross-validation fold, let $\mathcal{I}_{\text{train}}$ and $\mathcal{I}_{\text{test}}$ index the training and test participants, respectively.

We define

$$\mathbf{X}_{\text{train}} \in \mathbb{R}^{n\times v}, \quad \mathbf{X}_{\text{test}} \in \mathbb{R}^{m\times v},$$
$$\mathbf{C}_{\text{train}} = \mathbf{C}[\mathcal{I}_{\text{train}}, :] \in \mathbb{R}^{n\times k}, \quad \mathbf{C}_{\text{test}} = \mathbf{C}[\mathcal{I}_{\text{test}}, :] \in \mathbb{R}^{m\times k}.$$

Confounds were standardized using training-set statistics only. Let

$$\boldsymbol{\mu} = \tfrac{1}{n}\mathbf{1}_n^\top \mathbf{C}_{\text{train}}, \quad \boldsymbol{\sigma} = \text{sd}(\mathbf{C}_{\text{train}}) \in \mathbb{R}^k.$$

The standardized confound matrices are then

$$\widetilde{\mathbf{C}}_{\text{train}} = (\mathbf{C}_{\text{train}} - \mathbf{1}_n\boldsymbol{\mu}^\top) \oslash \boldsymbol{\sigma}, \quad \widetilde{\mathbf{C}}_{\text{test}} = (\mathbf{C}_{\text{test}} - \mathbf{1}_m\boldsymbol{\mu}^\top) \oslash \boldsymbol{\sigma}.$$

An intercept was added to each design matrix:

$$\widehat{\mathbf{C}}_{\text{train}} = [\mathbf{1}_n \;\; \widetilde{\mathbf{C}}_{\text{train}}] \in \mathbb{R}^{n\times(k+1)}, \quad \widehat{\mathbf{C}}_{\text{test}} = [\mathbf{1}_m \;\; \widetilde{\mathbf{C}}_{\text{test}}] \in \mathbb{R}^{m\times(k+1)}.$$

Regression coefficients were estimated on the training split using ordinary least squares:

$$\boldsymbol{\beta} = (\widehat{\mathbf{C}}_{\text{train}}^\top \widehat{\mathbf{C}}_{\text{train}})^{-1} \widehat{\mathbf{C}}_{\text{train}}^\top \mathbf{X}_{\text{train}}. \tag{5}$$

Residualized feature matrices were obtained as

$$\mathbf{X}_{\text{train}}^{\text{resid}} = \mathbf{X}_{\text{train}} - \widehat{\mathbf{C}}_{\text{train}}\boldsymbol{\beta}, \quad \mathbf{X}_{\text{test}}^{\text{resid}} = \mathbf{X}_{\text{test}} - \widehat{\mathbf{C}}_{\text{test}}\boldsymbol{\beta}.$$

These residualized features were subsequently standardized using training-set statistics and used as inputs to all multivariate decoding analyses.

#### Permutation-Based Cluster-Level Inference for MVPA

**Algorithm 1** Cluster-Level Correction for MVPA using Permutation Testing

**Require:** Observed accuracy map $\mathbf{M}$, permutation maps $\mathbf{P}$, voxelwise threshold $p_{\text{thresh}}$, minimum cluster size $k$, region mask $\mathbf{R}$, voxel volume $v^3$, number of permutations $N$

Load analysis mask $\mathbf{B}$ and extract voxel indices $\mathcal{I} = \text{where}(\mathbf{B} > 0)$
Compute voxelwise null distributions from $\mathbf{P}$ using one-sided testing
Compute voxelwise percentile threshold map $\mathbf{T} = \text{Percentile}(\mathbf{P}, 1 - p_{\text{thresh}})$
Threshold observed map: $\mathbf{M}_{\text{thresh}} = \mathbf{M} > \mathbf{T}$
Identify connected clusters in $\mathbf{M}_{\text{thresh}}$ within mask $\mathbf{R}$ with size $> k \cdot v^3$
**for** each observed cluster $c$ **do**
    Compute cluster statistic $s_c = \sum_{i \in c}(\mathbf{M}_i - \mathbf{T}_i)$
**end for**
**for** $p = 1$ to $N$ **do**
    Compute threshold $\mathbf{T}^{(p)}$ from $\mathbf{P}^{(p)}$
    Identify clusters in $\mathbf{P}^{(p)} > \mathbf{T}^{(p)}$
    Compute maximum cluster statistic $s_p^{\max}$
**end for**
Form null distribution $\{s_1^{\max}, \dots, s_N^{\max}\}$
**for** each $s_c$ **do**
    Compute FWER-corrected $p$-value: $p_c = \frac{1}{N} \sum_{p=1}^{N} \mathbb{I}[s_c \leq s_p^{\max}]$
**end for**
**return** Significant clusters and corrected $p$-values

#### fMRI Modelling

To ensure transparency and reproducibility of all fMRI analyses, we provide a detailed overview of the modeling framework used at the preprocessing, first-level, and group-level stages. Across all analyses, we made modeling choices to minimize confounding, avoid information leakage, and enable interpretable inference at each level.

We preprocess functional MRI data using fMRIPrep (v23.1.3), followed by signal cleaning and confound regression implemented in Nilearn (v0.10.0). Preprocessing included correction for B0 field inhomogeneities, intensity non-uniformity correction, skull stripping, tissue segmentation, surface reconstruction, spatial normalization, head motion correction, slice-timing correction, and resampling to standard space. A comprehensive set of motion-, physiological-, and noise-related confounds was estimated, including rigid-body motion parameters and their derivatives, framewise displacement, DVARS, global signal, white matter and cerebrospinal fluid signals, and CompCor-derived components. High-dimensional confound sets were standardized and reduced using principal component analysis (PCA; 95% variance explained) before regression. Signal cleaning included detrending, high-pass filtering (200 s cut-off), confound regression, and standardization, after which we concatenated run-wise time series at the participant level.

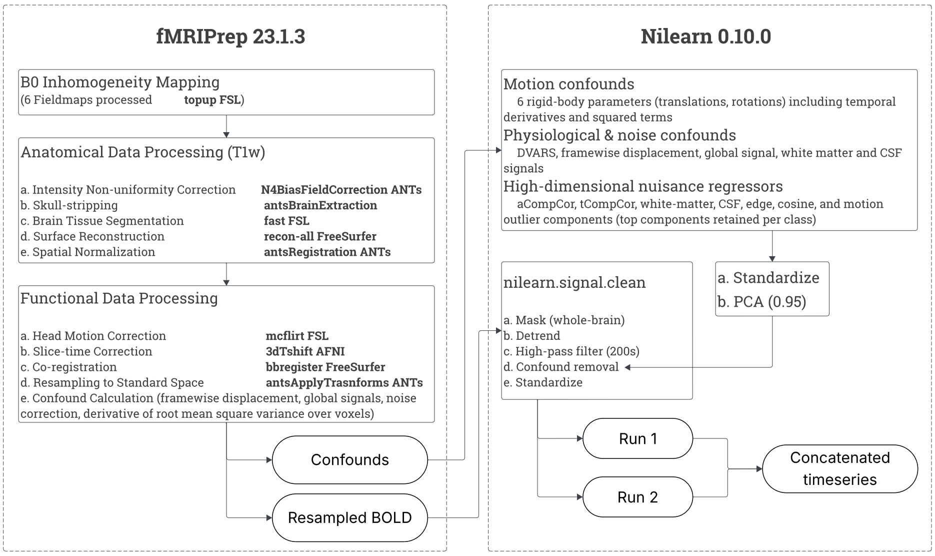


**Fig. S10**: **fMRI preprocessing and signal cleaning pipeline.** Overview of preprocessing and signal cleaning steps applied to functional MRI data. Left: fMRIPrep workflow, including B0 inhomogeneity correction, anatomical preprocessing (bias-field correction, skull stripping, tissue segmentation, surface reconstruction, spatial normalization), and functional preprocessing (motion correction, slice-timing correction, co-registration, resampling, and confound estimation). Right: Nilearn-based confound regression and signal cleaning, including regression of motion, physiological, and noise confounds, high-pass filtering (200 s cutoff), detrending, and standardization. High-dimensional confound sets were standardized and reduced using PCA (95% variance explained). We cleaned and concatenated the across runs at the participant level for subsequent analyses.

At the first (within-participant) level, we modeled the task-evoked activity during the training phase using a general linear model (GLM) comprising regressors capturing both sustained block-level engagement and linear changes in behavior across blocks. Specifically, separate regressors modeled the mean block response and the linear trend within blocks, each convolved with a canonical SPM hemodynamic response function and augmented with temporal and dispersion derivatives to account for variability in response timing and shape. Figure S11 displays the exact first-level design matrix used for estimation, along with the corresponding contrast vectors used to test mean block-related activity and linear block-related activity independently.

At the group level, we incorporated subject-wise summary measures of behavioral performance, task structure, and motor control, as well as questionnaire-based trait measures, into a second-level design matrix. This matrix included demographic covariates, task counterbalancing variables, technical control variables, and psychological measures, along with an intercept term. Figure S12 shows the full group-level

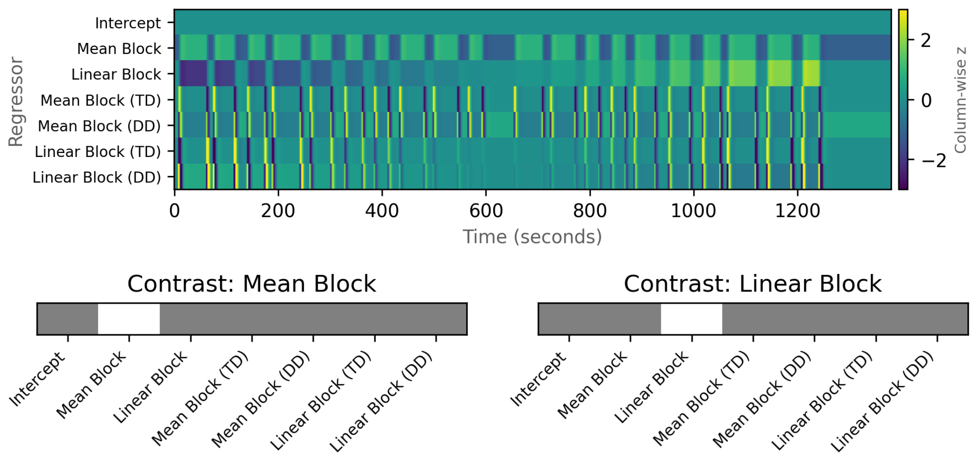


**Fig. S11**: **First-level design matrix and contrasts used for training-phase GLM analyses.** Top: Exact first-level design matrix (regressors × time) used to model training-phase BOLD responses. Regressors include an intercept, sustained mean block regressor, linear block regressor, and corresponding temporal (TD) and dispersion (DD) derivatives. Values are displayed as column-wise z-scores for visualization (clipped), while model fitting used the original regressors. Bottom: Contrast vectors used to test mean block-related activity and linear block-related activity, respectively. We include intercept and derivative regressors in contrasts of interest.

design matrix and the correlation structure among regressors, providing a comprehensive view of the statistical dependencies in the model. All group-level analyses tested effects of interest using contrasts that assigned unit weight to the regressor of interest and zero weight to all other regressors, including the intercept.

Together, these figures provide a complete, explicit account of the modeling pipeline used for fMRI analyses at all stages, from preprocessing through group-level inference.

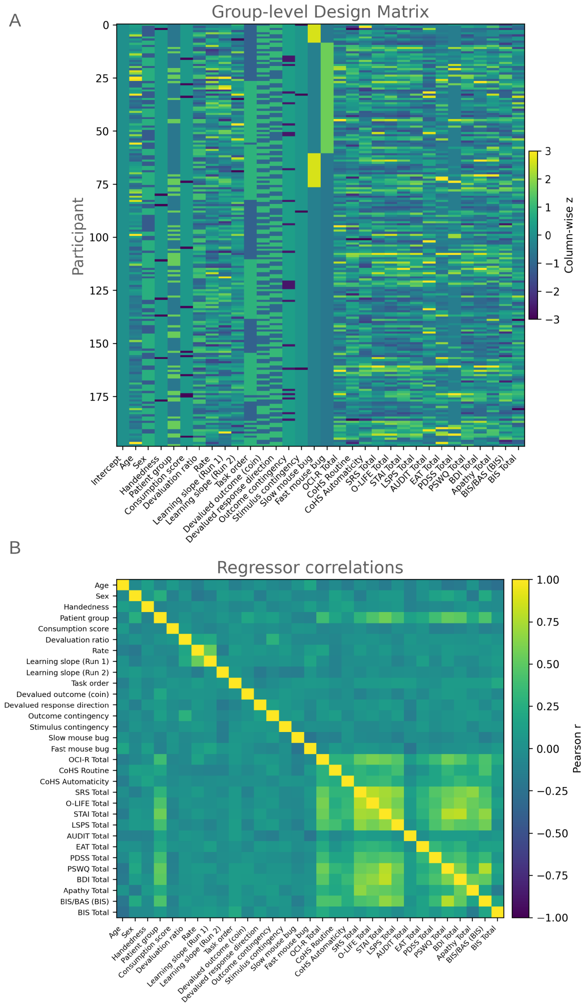


**Fig. S12**: **Group-level design matrix and regressor correlations.** **(A)** Group-level design matrix (participants × regressors) used for second-level analyses, including demographic variables, behavioral measures, task counterbalancing variables, technical control variables, and questionnaire-based trait measures. Values are shown as column-wise z-scores for visualization only. **(B)** Pearson correlation matrix across group-level regressors, illustrating the dependency structure among predictors included in the model. This figure provides a comprehensive view of the statistical structure underlying group-level inference.
